# Vascular smooth muscle cell loss, but not neuroinflammation, drives cerebrovascular reactivity impairment in Alzheimer’s disease

**DOI:** 10.64898/2025.12.01.691642

**Authors:** Xiuli Yang, Yuguo Li, Minmin Yao, Adnan Bibic, Wenzhen Duan, Hanzhang Lu, Zhiliang Wei

## Abstract

**INTRODUCTION:** Cerebrovascular reactivity (CVR) impairment is a key feature of Alzheimer’s disease (AD), but its mechanistic basis remains unclear. This study examined whether vascular smooth muscle cell (VSMC) loss, rather than amyloidosis or neuroinflammation, underlies CVR deficits.

**METHODS:** Non-contrast MRI, including phase-contrast and pseudo-continuous arterial spin labeling, was performed in mouse models of amyloidosis (5xFAD), VSMC degeneration (CADASIL), and lipopolysaccharide-induced neuroinflammation. Characterization of vascular, amyloid-β, and inflammatory markers were performed for pathological assessment.

**RESULTS:** CVR impairment emerged only when VSMC loss was present in CADASIL mice and at older ages in 5xFAD mice (9–12 months). Amyloid-β deposition occurred earlier than VSMC loss or CVR decline. Neuroinflammation primarily altered baseline cerebral blood flow without affecting CVR or VSMC integrity.

**DISCUSSION:** These findings identify VSMC degeneration as an important driver of CVR impairment independent of cerebral amyloid angiopathy or inflammation, highlighting vascular integrity as a potential therapeutic target in AD.

**Highlights:**

- Cerebrovascular reactivity (CVR) impairment occurred in 5xFAD mice only when vascular smooth muscle cell (VSMC) loss was present
- 5xFAD mice exhibited prominent parenchymal but minimal vascular amyloid-β deposition
- VSMC developmental deficiency resulted in CVR impairment in a small-vessel disease (SVD) model
- Neuroinflammation primarily altered baseline cerebral blood flow (CBF) without affecting CVR

## 1. INTRODUCTION

Cerebrovascular reactivity (CVR) reflects the capacity of cerebral blood vessels to dilate or constrict in response to vasoactive stimuli, thereby maintaining an adequate supply of oxygen and nutrients to brain tissue in a dynamic manner subject to the neuronal activity.^1^ This capacity is crucial because the brain has very high metabolic demand (∼2% of body weight but occupied ∼20% of the total energy^2^) however almost no energy storage.^3^ Baseline cerebral blood flow (CBF) reflects the resting supply capacity of the microvasculature,^4^ whereas cerebrovascular reactivity (CVR) gauges the reserve ability of the vascular network to adapt to changing metabolic demands.^5^ Together, they embody the two pillars of microvascular physiology — supply and reserve. Because the vascular reserve can be consumed to maintain perfusion under stress, CVR impairment may occur even when resting CBF appears normal and CVR often correlates more strongly with cognitive performance than resting CBF.^6^

Deficits in CVR predispose the brain to hypoxia and metabolic strain, accelerating neurodegenerative processes.^7,8^ In both clinical and preclinical contexts, CVR is increasingly recognized as a potential early biomarker of vascular dysfunction that precedes overt structural changes or cognitive decline.^9^ Age-related attenuation of CVR is widely reported across cross-sectional and longitudinal studies, even in cognitively normal adults without noticeable vascular disease.^10-12^ In older adults with cognitive impairment, whole-brain CVR tends to be lower compared with cognitively normal peers, and higher CVR is associated with better cognitive performance.^13^ CVR deficits are further confirmed in asymptomatic and prodromal Alzheimer’s disease (AD) patients.^14^ In cerebral small vessel disease (SVD), CVR impairment is linked with worse cognitive performance and greater concurrent neuroimaging burden.^15^ Beyond single-site reports, impaired CVR has been shown across multiple cohorts to associate with global cognitive metrics (e.g. the Montreal Cognitive Assessment, MoCA) and executive function in older adults.^16^ These converging findings argue for incorporating CVR into clinical workflows. Clinical vascular-targeted therapies intended to improve cerebrovascular function can be evaluated by whether they improve CVR — making it a potential outcome metric.^17^ Meanwhile, CVR can stratify cohorts to identify high-risk individuals and likely responders, facilitating precision medicine across preclinical and clinical studies—for example, assessing vascular risk in AD patients treated with recently approved anti-amyloid monoclonal antibodies^18,19^.

Despite growing evidence in humans linking CVR to cognitive impairment, small vessel disease burden, and risk stratification outcomes, mechanistic delineation of the vascular drivers of CVR decline remains limited. Cerebral amyloid angiopathy (CAA) has been implicated as a contributor to microvascular dysfunctions, including disease related CBF declines^20^ and CVR impairment^21^. However, whether substantial CAA is required for CVR deficits or whether neuroinflammation, a pervasive feature of neurological disease, modulates CVR remains unclear. Here, we integrate non-invasive MRI with pathological characterizations across mouse models of AD, SVD, and neuroinflammation to dissect the vascular determinants of CVR. We show that loss of vascular smooth-muscle cells (VMSC), rather than CAA or inflammation, underlies CVR impairment. These findings refine the interpretation of human CVR alterations, highlight VSMC integrity as a key regulator of cerebrovascular function, and establish a potential translational framework for linking microvascular pathology to functional vascular imaging biomarkers.

## 2. METHODS

### 2.1 General procedures

The experimental protocols for this study were approved by the Johns Hopkins Medical Institution Animal Care and Use Committee and conducted in accordance with the National Institutes of Health guidelines for the care and use of laboratory animals. Data reporting complied with the ARRIVE 2.0 guidelines. A total number of N = 163 C57BL/6 mice (age: 4–20 months; body weight: 20–50 grams; 78 females [F], 85 males [M]) were used in this study. All mice were housed in a quiet environment with a 12-h day/night cycle and had ad libitum access to food and water.

### 2.2 Mouse models

Three mouse models were used: one AD model, one SVD model, and one neuroinflammation model. The 5xFAD line (Jackson Laboratory, stock #034848) is an APP/PS1 double-transgenic model carrying five familial AD mutations: the Swedish (K670N/M671L), Florida (I716V), and London (V717I) mutations in APP, and the M146L and L286V mutations in PSEN1.^22^ Autosomal-dominant arteriopathy with subcortical infarcts and leukoencephalopathy (CADASIL) is a transgenic SVD model conditionally expressing a mutant NOTCH3 allele, leading to developmental deficiency of vascular smooth-muscle cells.^23^ CADASIL mice were obtained by cross breeding the 129-Gt(ROSA)26Sor^tm2(NOTCH3*C455R)Sat^Mmjax line (Mutant Mouse Resource and Research Center, stock # 033000-JAX) with the SM22-Cre driver line (Jackson Laboratory, stock 017491).^23^ All described mouse lines were secured from The Jackson Laboratory and maintained through local breeding colonies to reach the targeted sample sizes for the experiments. Genotyping was performed twice, initially prior to group assignment and subsequently after euthanasia, to ensure scientific rigor. Neuroinflammation was induced by intraperitoneal lipopolysaccharide (LPS) injection (2 mg/kg) in C57BL/6 mice.^24^

### 2.3 MRI experiments

All MRI experiments were performed on an 11.7T Bruker Biospec system (Bruker, Ettlingen, Germany) with a horizontal bore and an actively shielded pulse-field gradient (maximum intensity of 0.74 T/m). Images were acquired using a 72-mm quadrature volume resonator as the transmitter, and a four-element (2×2) phased-array coil as the receiver. Magnetic field homogeneity over the mouse brain was optimized using global shimming (up to the second order) based on a pre-acquired subject-specific field map. Inhalational isoflurane, delivered with medical air (21% O2 and 78% N2) or hypercapnia gas (5% CO2, 21% O2, and 74% N2) at a flow rate of 0.75 L/min, was used to minimize stress and motion in the mice. Anesthesia induction was achieved with 1.5-2.0% isoflurane for 15 minutes. At the 10^th^ minute under induction, the mouse was placed onto a water-heated animal bed with temperature control and positioned with a bite bar, a pair of ear pins, and a custom-built 3D-printed holder before entering the magnet.^25^ After induction, the isoflurane concentration was reduced to 1.0% for maintenance during the MRI scans. Respiration was observed during the experiments using an MRI-compatible monitoring system (SA Instruments, Stony Brook, NY, USA). Isoflurane dose was adjusted as needed to maintain a respiration rate of 70–120 breaths per minute. Experiments were terminated if respiration rates dropped below 50 breaths per minute for more than 2 minutes.

Mice were scanned in a randomized order, following a previously reported scheme.^26^ Briefly, mice were preassigned consecutive numbers starting from one. A set of pseudorandom numbers was generated using the MATLAB (MathWorks, Natick, MA, USA) “rand” function, and the ranks of these numbers (from largest to smallest) determined the experimental order.

#### Vascular and metabolic responses to hypercapnia challenge (*N* = 58; 22F/36M)

A 5% CO2 hypercapnia challenge^5^ was used to probe vascular and metabolic responses. Dynamic phase-contrast (PC) MRI was acquired over 5 min, with hypercapnia initiated at 0.3 min (*n* = 10; 5 females [F]/5 males [M]). Acquisition parameters followed a published PC MRI protocol:^27,28^ TR/TE = 15/3.2 ms; field of view (FOV) = 15 × 15 mm²; in-plane resolution = 50 × 50 µm²; slice thickness = 0.5 mm; velocity encoding (VENC) = 20 cm/s; number of averages = 2; dummy scans = 8; receiver bandwidth = 100 kHz; flip angle = 25°; partial-Fourier factor = 0.7; and temporal resolution = 0.3 min (18 s).

In a separate session, PC and T2-relaxation-under-spin-tagging (TRUST)^29^ MRI were performed under medical air followed by 5% CO2, separated by a 5-min transition, to assess hypercapnia-induced metabolic modulation (*n* = 12 [6F/6M]). PC MRI targeted the major cerebral feeding arteries—the left and right internal carotid arteries and the basilar artery.^30^ Key parameters for TRUST MRI were:^28^ TR/TE = 3500/4.9 ms; FOV = 16 × 16 mm^2^; in-plane resolution = 125 × 125 µm²; slice thickness = 0.5 mm, inversion slab thickness = 2.5 mm, post-labeling delay = 1000 ms, effective TE (eTE) = 0.3/20/40 ms, echo spacing of eTE = 5.0 ms, and scan duration = 2.8 min with an 8-segment gradient-echo echo-planar imaging (EPI) acquisition. In addition, a fast spin-echo (FSE) scan was performed to estimate brain volume using a reported protocol.^31^

To examine the anesthetic effect on CVR, PC MRI was performed under medical air and then 5% CO2, separated by a 5-min transition, in a cohort of 36 mice (11F/25M). In a subgroup of 10 mice (5F/5M), CVR measurements were repeated consecutively three times within the same session, yielding a total of 56 data points.

Processing of PC datasets followed published procedures.^27,28^ Briefly, a region of interest (ROI) was manually delineated on the complex-difference magnitude image and transferred to the phase-derived velocity map. Volumetric flow for each artery was obtained by integrating through-plane velocity over the ROI cross-section, then summing across the feeding arteries. Total inflow was normalized by brain weight to yield CBF in the unit of ml/100g/min. CVR was calculated as 100·(CBF_post_-CBF_pre_)/CBF_pre_ with a unit of %.

#### CVR mapping in 5xFAD mice (*N* = 46; 26F/20M)

PC and pseudo-continuous arterial spin labeling (pCASL) MRI were used for evaluating global and regional CVR. 5xFAD mice at 9-12 months of age (*n* = 16, 7F/9M), WT mice at 9-12 months of age (*n* = 12, 9F/3M), 5xFAD at 5 months of age (*n* = 10, 5F/5M), and WT mice at 5 months of age (*n* = 8, 5F/3M) were examined for CVR.

A two-scan pCASL scheme developed to improve the labeling performance under magnetic-field inhomogeneity was used.^32^ Key parameters of pCASL MRI followed previous reports:^33^ TR/TE = 3000/11.8 ms; FOV = 15 × 15 mm²; in-plane resolution = 157 × 157 µm^2^; slice thickness = 0.75 mm; interslice gap = 0.25 mm; number of slices = 12; receiver bandwidth = 300 kHz; labeling duration = 1800 ms; inter-labeling pulse delay = 1.0 ms; labeling pulse width = 0.4 ms; labeling plane thickness = 1.0 mm; labeling pulse flip angle = 40°; mean gradient = 0.01 T/m (label scans) and 0 T/m (control scans); post-labeling delay = 300 ms; signal average = 25; and scan duration = 5.0 min using a two-segment spin-echo EPI acquisition.

Processing of pCASL data followed established procedures.^26^ Pairwise subtraction between control and labeled images was first performed to yield a difference image, which was then divided by an *M*_0_ image (obtained by scaling the control image^32^) to provide a perfusion index image. The perfusion index maps were co-registered and normalized to a mouse brain template,^34^ resized to recover the original acquisition resolutions, and rescaled by reference to the global CBF values (from PC MRI) to obtain absolute values. ROIs were drawn on the averaged control images to encompass the midbrain, isocortex, hippocampus, thalamus, hypothalamus, and striatum by reference to the Allen mouse brain atlas (https://atlas.brain-map.org/). Voxel-wise CBF values within each ROI were averaged to estimate the corresponding perfusion levels.

#### CVR mapping in CADASIL mice (*N* = 36; 18F/18M)

CADASIL mice at 9 months of age (*n* = 6; 3F/3M), WT mice at 9 months of age (*n* = 6; 3F/3M), CADASIL mice at 13 months of age (*n* = 12; 6F/6M), and WT mice at 13 months of age (*n* = 12; 6F/6M) were examined with PC and pCASL MRI for CVR mapping.

#### CVR mapping in neuroinflammation model (*N* = 15; 8F/7M)

A cohort of 15 mice was randomly divided into two groups to receive LPS (*n* = 10; 5F/5M) or saline injection (*n* = 5; 3F/2M). Mice were scanned pre-injection and two days post-injection for CVR mapping.^24^ Two male mice that received the LPS injection died before the post-injection MRI, possibly due to the relatively higher LPS injection volume associated with their larger body weights.

### 2.4 Immunostaining

Brain sections were collected and processed following previous reports.^35,36^ Briefly, mice were perfused transcardially with PBS, followed by phosphate-buffered 4% paraformaldehyde. The brains were fixed in phosphate-buffered 4% paraformaldehyde for 24 h at 4 °C and dehydrated with 30% sucrose. Frozen brain tissue sections at the thickness of 30 μm were prepared using a cryostat. After the blocking procedure, the brain sections were incubated with primary antibodies for 2 days at 4 °C. Subsequently, sections were incubated for 2 hours with matched fluorescence-linked secondary antibodies (Jackson ImmunoResearch Laboratories, West Grove, USA). The primary antibodies used were Collagen IV (1:500, 2150-1470, Bio-Rad, Hercules, USA), alpha-smooth muscle actin (α-SMA) (1:250, C6198, Millipore Sigma, Burlington, USA), Iba1 (1:500, 019-19741, Fujifilm, Tokyo, Japan), CD68 (1:250, MCA1957, Bio-Rad, Hercules, USA), and 6E10 (1:500, 803001, BioLegend, San Diego, USA). More antibody details were summarized in Supporting Table S1. For each mouse, three slices from frontal and parietal regions at similar cerebral positions were selected and stained.^37^ Fluorescently labeled samples were imaged using a Zeiss LSM 780 FCS confocal microscope (Zeiss, Oberkochen, Germany). Representative slices from each group were initially inspected at higher magnification (×40 or ×63) to confirm staining quality, after which all slices were scanned at ×10 magnification for quantitative analysis.^37^

Among the scanned mice, a subgroup of 5xFAD mice at 5 months of age (*n* = 5 [3F/2M] WT, *n* = 5 [3F/2M] 5xFAD) and 9–12 months of age (*n* = 5 [3F/2M] WT, *n* = 5 [1F/4M] 5xFAD) was examined using co-staining for Collagen IV and α-SMA, Collagen IV and 6E10, and Iba1 and CD68. A subgroup of CADASIL mice at 9 months (*n* = 5 [3F/2M] WT, *n* = 6 [3F/3M] CADASIL) and 13 months (*n* = 6 [3F/3M] WT, *n* = 6 [3F/3M] CADASIL) of age was examined using co-staining for Collagen IV and α-SMA. A subgroup of neuroinflammation-model mice (*n* = 5 [3F/2M] post-saline, *n* = 5 [3F/2M] post-LPS) was examined using co-staining for Collagen IV and α-SMA, and Iba1 and CD68. A separate cohort of C57BL/6 mice (*n* = 5, 2F/3M) without injection was included as a pre-injection reference for the neuroinflammation model.

### 2.5 Transmission electron microscopy (N = 3; 2F/1M)

Transmission electron microscopy (TEM) was performed to examine the presence of granular osmiophilic material (GOM), a characteristic structural hallmark of CADASIL. Three mice at 9 months of age (*n* = 1 [1F] WT; *n* = 2 [1F/1M] CADASIL) were examined. Brain tissues containing different regions were processed using standard fixation and resin-embedding procedures. Ultrathin sections (60–70 nm) were cut using a Leica UC7 ultramicrotome, collected on copper grids, and stained with UranyLess and lead citrate. TEM images were acquired using a Thermo Fisher Talos L120C operating at 120 kV and equipped with a Ceta CMOS camera.

### 2.6 Data processing

MRI data were processed with custom MATLAB scripts (MathWorks, Natick, MA, USA). Processing of microscope images was performed as previously described.^38^

### 2.7 Statistical analyses

Group-wise comparisons were performed using the Student’s *t*-test, with *t*-statistics and associated degrees of freedom reported. Linear regression and linear mixed-effects models were used to examine alterations in MRI- and histology-derived measurements. Models were fitted using restricted maximum likelihood estimation, and degrees of freedom were determined by the residual method. Effect estimates (β) and corresponding *p*-values were reported. A threshold of *p* < 0.05 was considered statistically significant. Statistical details for all comparisons performed in this study were summarized in Supporting Table S2.

## 3. RESULTS

### 3.1 Hypercapnia challenge modulates vascular function without affecting metabolism in mice

To standardize the hypercapnia challenge protocol in mice, we first examined CBF dynamics using PC MRI across a 5-min transition (Fig. 1a). An increase in CBF following hypercapnia was evident from the progressive rise in blood-flow velocity at 0.3, 1.8, 3.3, and 4.8 min post-onset (Fig. 1b). At the group level (*n* = 10), the percent change in CBF exhibited a biphasic pattern (Fig. 1c) that was modeled by y = 40.53[1-e^-(x-1.35)/1.21^]·H(x-1.35), where H(x) denotes the Heaviside function. The initial flat period (≤1.35 min) reflects the delay required for 5% CO2 to travel through the delivery tubing and the cardiopulmonary system before taking effect, after which a rapid increase in CBF indicates the vasodilatory response to CO2. According to the fitted model, the 5-min transition achieved 95.1% of the maximal CBF change and was considered adequate for observing the vascular responses to hypercapnia. Intravenous acetazolamide administration,^39^ an alternative approach to assess vascular reactivity, elicits a similar response by inhibiting carbonic anhydrase and thus indirectly increasing blood CO2 through accumulating metabolism-generated CO2. When defining the buildup time as the duration needed to reach 63.21% of the final CBF change (i.e., 1 - *e*^-1^), hypercapnia (1.21 min) was more efficient than acetazolamide (1.62 min), consistent with the fact that inhaled CO2 directly elevates blood CO2 level with a high partial pressure. Given the simpler implementation of gas delivery compared with intravenous catheterization, hypercapnia is more suitable for large-scale studies. Therefore, in subsequent studies, hypercapnia with a 5-min transition period was consistently employed alongside PC and pCASL MRI (Figure S1a) for CVR assessment.

**Figure 1.**
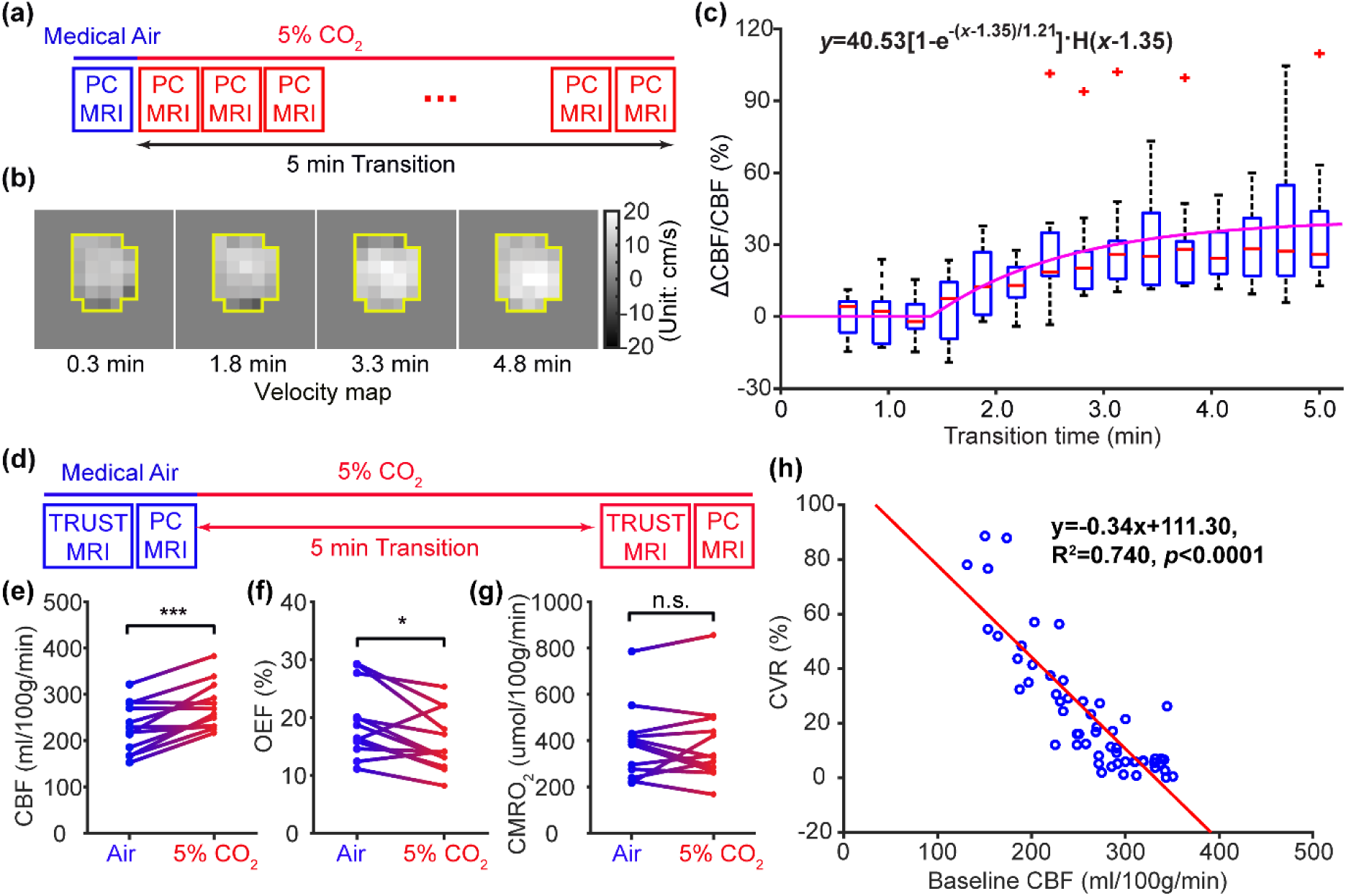
Vascular and metabolic responses to hypercapnia. Data were collected from N = 58 mice (22 females [F], 36 males [M]). (a) Schematic diagram of the dynamic phase-contrast (PC) MRI over a 5-min transition (*n* = 10; 5F/5M). (b) Representative PC-derived velocity maps at 0.3, 1.8, 3.3, and 4.8 min. (c) Percent change in cerebral blood flow (CBF) relative to pre-hypercapnia baseline (*n* = 10; 5F/5M). (d) Schematic of metabolic measurements using T2-relaxation-under-spin-tagging (TRUST) MRI. (e, f, and g) Comparison of CBF, oxygen extraction fraction (OEF), and cerebral metabolic rate of oxygen (CMRO2) between air and 5% CO2 conditions (*n* = 12; 6F/6M). For group-wise comparisons: \**p* < 0.05; \*\*\**p* < 0.001; n.s., not significant. (h) Correlation between CBF and cerebrovascular reactivity (CVR) (*n* = 36; 11F/25M).

TRUST and PC MRI were combined to assess metabolic responses to hypercapnia across a 5-min transition (*n* = 12; Fig. 1d). CBF increased significantly after hypercapnia (*p* < 0.001; full statistics available in Supporting Table S2; Fig. 1e), accompanied by a significant reduction in oxygen extraction fraction (OEF) (*p* < 0.05; Fig. 1f). Using the Fick principle, the cerebral metabolic rate of oxygen (CMRO2) was estimated and remained unchanged (*p* > 0.05; Fig. 1g). These results indicate that hypercapnia increases CBF without changing metabolism (isometabolic response). Thus, hypercapnia chiefly indexes vascular reactivity with minimal metabolic confounding in mice.

Isoflurane anesthesia is widely used in mouse neuroimaging to minimize stress and motion and increases CBF in a dose-dependent manner.^40,41^ Consequently, substantial inter-subject variation in CBF can arise from differences in anesthetic depth. We therefore examined the CBF-CVR relationship in C57BL/6 mice (*n* = 36). CVR was negatively correlated with CBF (y = -0.34x + 111.30; *R*² = 0.740, *p* < 0.0001; Fig. 1h), consistent with the partial saturation of vascular reserve by isoflurane. To account for the CBF-CVR correlation, CBF was included as a covariate in later regression models comparing CVR in disease models.

### 3.2 α-SMA loss is more closely associated with CVR impairment than amyloidosis

An age effect (*p* < 0.001) but no genotype effect was observed in the brain volume of 5xFAD model (Fig. 2a), consistent with reports that tauopathy, not amyloidosis, drives neuronal loss.^26^ Global CBF showed no significant genotype- or age-dependent differences (Fig. 2b), likely due to opposing regional CBF alterations across the midbrain (*p* = 0.005), isocortex (*p* = 0.045), and thalamus (*p* = 0.022) (Fig. S1b–g). A significant genotype × age interaction was observed in global CVR (*p* = 0.036; Fig. 2c). Referencing to the CBF maps (Fig. 2d), this interaction was also evident in the isocortical (*p* = 0.027) and hippocampal (*p* = 0.033) CVR values (Fig. 2e & Fig. S1h–k). Immunofluorescent images of three whole slices stained for α-SMA and Collagen IV were collected for each mouse (Fig. 2f; Fig. S2a) to enable regional analyses. Higher-magnification views of the hippocampus (Fig. 2f) and isocortex (Fig. S2b) confirmed comparable staining quality across regions. The α-SMA coverage index, defined as the ratio of the co-localized α-SMA– and Collagen IV-positive area (α-SMA^+^Col-IV^+^) to the total Collagen IV-positive area (Col-IV^+^), showed a significant genotype × age interaction in both the isocortex (*p* = 0.023) and hippocampus (*p* = 0.024) (Fig. 2g). The concurrent genotype × age interaction effects in CVR and α-SMA coverage suggest a potential mechanistic link between CVR and VSMC integrity.

**Figure 2.**
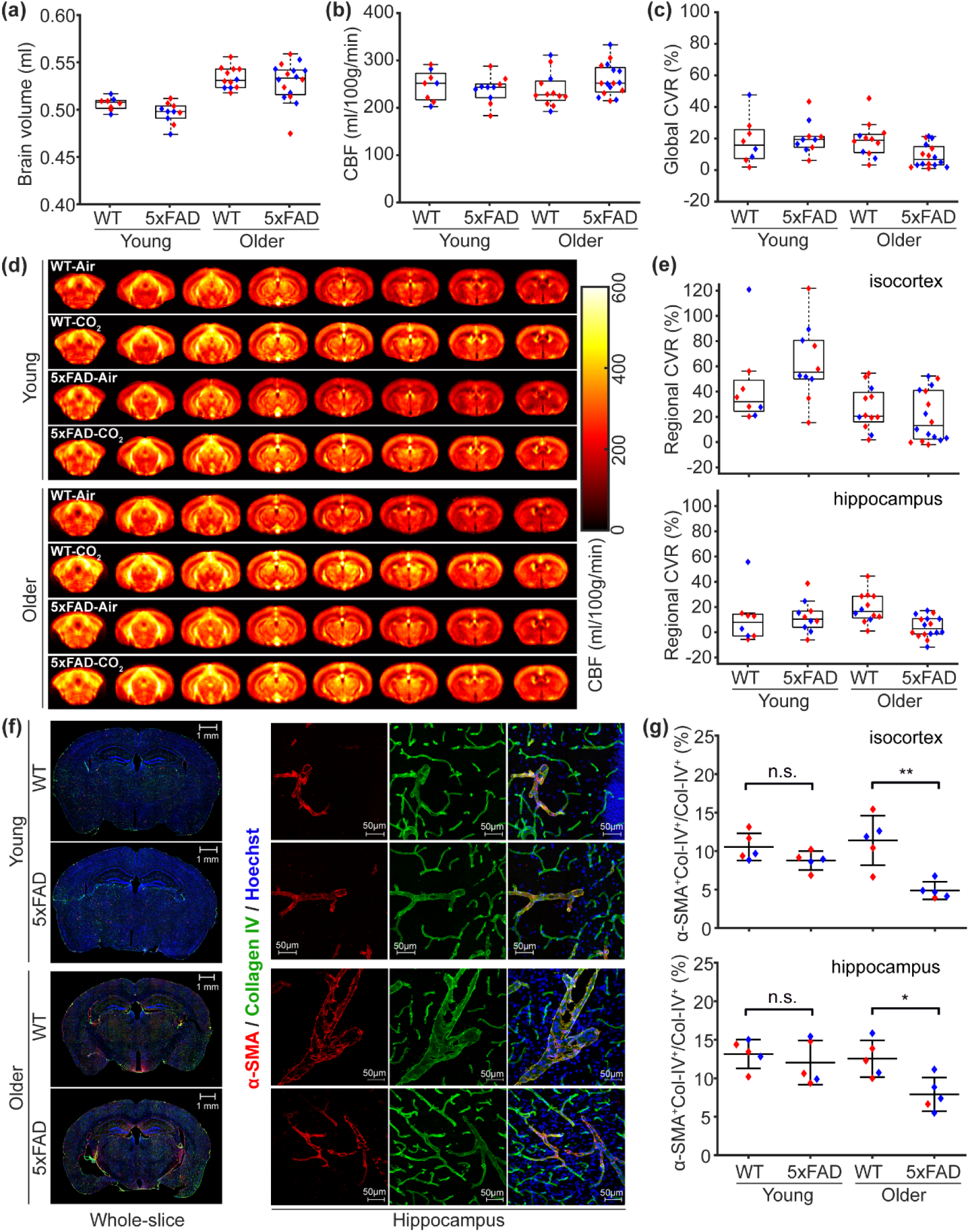
Characterization of the 5xFAD model. Comparisons of MRI measurements across young wild-type (WT; *n* = 8, 5F/3M), young 5xFAD (*n* = 10, 5F/5M), older WT (*n* = 12, 9F/3M), and older 5xFAD (*n* = 16, 7F/9M) mice, including brain volume (a), cerebral blood flow (CBF; b), global cerebrovascular reactivity (CVR; c), regional mean CBF maps (d), and isocortical and hippocampal CVR (e). Red and blue dots denote female and male mice, respectively. (f) Representative immunofluorescent images of α-SMA and Collagen IV staining. Scale bars: 1 mm (whole-slice views) and 50 µm (regional views). (g) Comparisons of α-smooth muscle actin (α-SMA) coverage index across young WT (*n* = 5, 3F/2M), young 5xFAD (*n* = 5, 3F/2M), older WT (*n* = 5, 3F/2M), and older 5xFAD (*n* = 5, 1F/4M) mice, which is defined as the ratio of the co-localized α-SMA and Collagen IV-positive area (α-SMA^+^Col-IV^+^) to the total Collagen IV-positive area (Col-IV^+^). For group-wise comparisons: \**p* < 0.05; \*\**p* < 0.01; n.s., not significant.

5xFAD mice have been reported to exhibit substantial parenchymal but minimal vascular amyloid-β (Aβ) deposition.^22^ To confirm this, we performed co-staining of 6E10 and Collagen IV. At the whole-slice level, 5xFAD mice showed markedly higher Aβ deposition than age-matched WT controls (Fig. 3a). Higher-magnification views of the isocortex and hippocampus (Fig. 3b) displayed an age-dependent increase in Aβ burden. At the group level, the Aβ coverage index, defined as the ratio of 6E10-positive area (6E10^+^) to the total regional area, showed significant genotype × age interaction effects in both the isocortex (*p* = 0.001) and hippocampus (*p* < 0.001) (Fig. 3c). Furthermore, the vascular Aβ coverage index, defined as the ratio of co-localized 6E10-positive and Collagen IV-positive area (6E10^+^Col-IV^+^) to the total Collagen IV-positive area (Col-IV^+^), exhibited a significant age-dependent increase in 5xFAD mice (isocortex: *p* = 0.028; hippocampus: *p* < 0.001; Fig. 3d). Notably, vascular Aβ accounted for only approximately 0–3% of the total Aβ burden (Fig. 3d).

**Figure 3.**
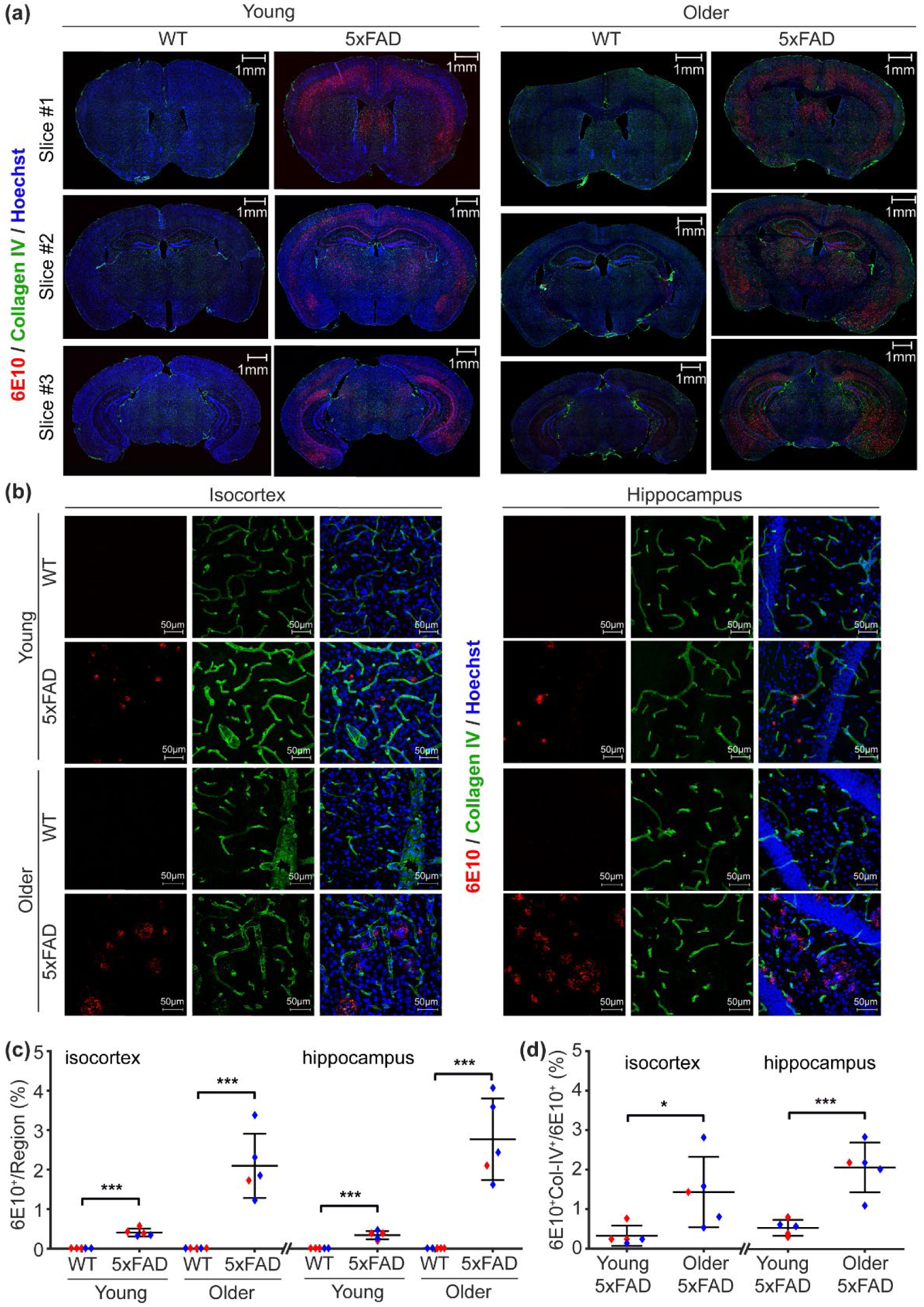
Amyloid-β deposition in the 5xFAD model. (a, b) Slice-by-slice and regional comparisons of immunofluorescent staining for amyloid-β (Aβ) in representative young wild-type (WT), young 5xFAD, older WT, and older 5xFAD mice. Scale bars: 1 mm (whole-slice views) and 50 µm (regional views). (c) Amyloid-β coverage index, defined as the ratio of 6E10-positive area (6E10^+^) to the total regional area, in the isocortex and hippocampus across the four groups. Red and blue dots represent female and male mice, respectively. Error bars indicate standard deviation. (d) Vascular amyloid coverage index, defined as the ratio of co-localized 6E10- and Collagen IV-positive area (6E10^+^Col-IV^+^) to the total 6E10-positive area (6E10^+^), in the isocortex and hippocampus. For group-wise comparisons: \**p* < 0.05; \*\*\**p* < 0.001.

In the hippocampus, significant reductions in α-SMA coverage and CVR were evident only in older 5xFAD mice (α-SMA coverage: *p* = 0.012; CVR: *p* < 0.001), whereas Aβ coverage was already elevated in both young (*p* < 0.001) and older (*p* < 0.001) cohorts. These results suggest that Aβ deposition occurs earlier in the pathological cascade, while VSMC degeneration is more directly linked to CVR impairment than to amyloid burden itself.

### 3.3 VSMC loss is sufficient to drive CVR impairment

A significant sex effect, but no genotype or age effect, was observed in brain volume in CADASIL mice (Fig. 4a), possibly reflecting sex-specific differences in brain development of this model. The absence of an age effect may be related to the relatively narrow observation window (i.e., 4 months). Both global and regional CBF values did not differ between CADASIL and WT mice (Fig. 4b; Fig. S3a–f). In contrast, global CVR exhibited significant genotype (*p* = 0.024), age (*p* = 0.004), and sex (*p* = 0.007) effects (Fig. 4c). According to the regional CBF maps (Fig. 4d), CVR impairment was most prominent in deep-brain regions, including the thalamus (*p* = 0.026; Fig. 4e), hypothalamus (*p* = 0.028; Fig. S3i), and striatum (*p* = 0.030; Fig. S3j), but not in the hippocampus (Fig. 4e) or other regions (Fig. S3g-h). The age-dependent increase in CVR across multiple regions (Session 3, Table S2) may indicate the emergence of compensatory mechanisms, while the sex effect on CVR likely reflects developmental differences, in agreement with the observed sex-related variation in brain volume. The genotype effect on CVR aligns with the known phenotype of CADASIL, characterized by VSMC developmental deficiency and subsequent VSMC loss.^23^

**Figure 4.**
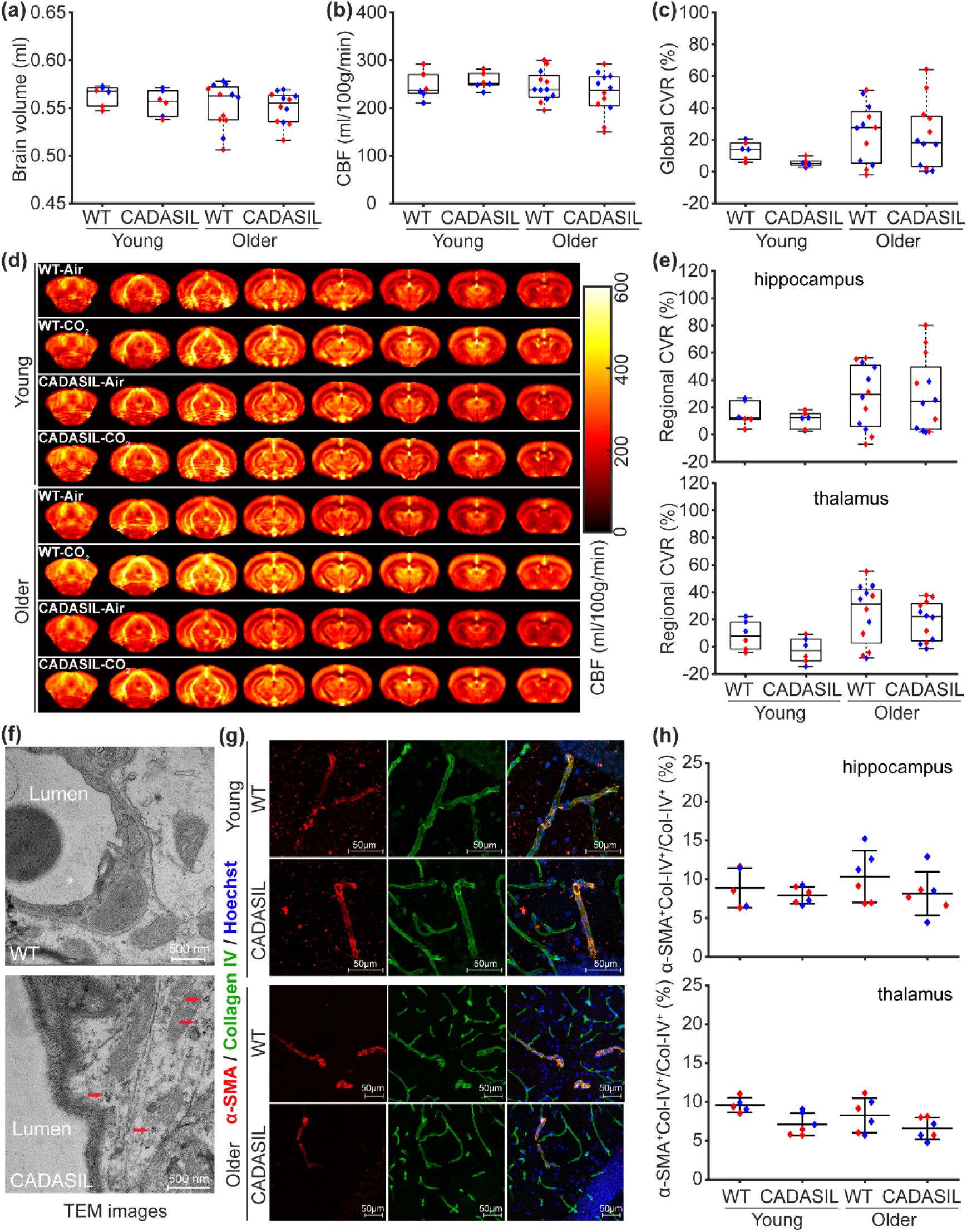
Characterization of cerebral autosomal dominant arteriopathy with subcortical infarcts and leukoencephalopathy (CADASIL) mice. Comparisons of MRI measurements across young wild-type (WT; *n* = 6, 3F/3M), young CADASIL (*n* = 6, 3F/3M), older WT (*n* = 12, 6F/6M), and older CADASIL (*n* = 12, 6F/6M) mice, including brain volume (a), cerebral blood flow (CBF; b), global cerebrovascular reactivity (CVR; c), regional mean CBF maps (d), and hippocampal and thalamic CVR (e). Red and blue dots denote female and male mice, respectively. (f) Transmission electron microscopy (TEM) images from representative young WT and CADASIL mice. (g) Representative immunofluorescent images showing α-smooth muscle actin (α-SMA, red) and Collagen IV (Col-IV, green) staining in the hippocampus. Scale bars: 50 µm. (h) Comparisons of the α-SMA coverage index across young WT (*n* = 5, 3F/2M), young CADASIL (*n* = 6, 3F/3M), older WT (*n* = 6, 3F/3M), and older CADASIL (*n* = 6, 3F/3M) mice. The α-SMA coverage index is defined as the ratio of the co-localized α-SMA- and Collagen IV-positive area (α-SMA^+^Col-IV^+^) to the total Collagen IV-positive area (Col-IV^+^).

TEM further confirmed the CADASIL pathology, revealing GOM consistently in 9-month-old CADASIL mice (red arrows; Fig. 4f). Immunofluorescent staining for α-SMA and Collagen IV demonstrated consistent staining quality across groups, as shown in higher-magnification images (Fig. 4g; Fig. S4). At the group level, α-SMA coverage showed a significant genotype effect (*p* = 0.008) in the thalamus but not in the hippocampus (Fig. 4h), consistent with the regional CVR pattern.

Collectively, these MRI and histological findings indicate that VSMC loss alone is sufficient to drive CVR impairment, independent of amyloidosis.

### 3.4 Neuroinflammation alters baseline CBF without affecting CVR

Brain volume was not affected by the injections (Fig. 5a). Global CBF decreased following LPS injection (*p* = 0.039; Fig. 5b), whereas global CVR was unaffected by injection or sex (Fig. 5c). Regional CBF maps allowed further spatial assessment (Fig. 5d). CBF was preserved in the hippocampus (Fig. 5e), midbrain (Fig. S5a), isocortex (Fig. S5b) but was significantly reduced in the thalamus (*p* = 0.018; Fig. S5c), hypothalamus (*p* = 0.035; Fig. S5d), and striatum (*p* = 0.017; Fig. S5e). In addition to the injection effect, significant sex-related differences were observed in both global and regional CBF (*p* ≤ 0.005; Session 4, Table S2). In contrast, CVR remained unchanged across all examined regions (Fig. S5f-j).

**Figure 5.**
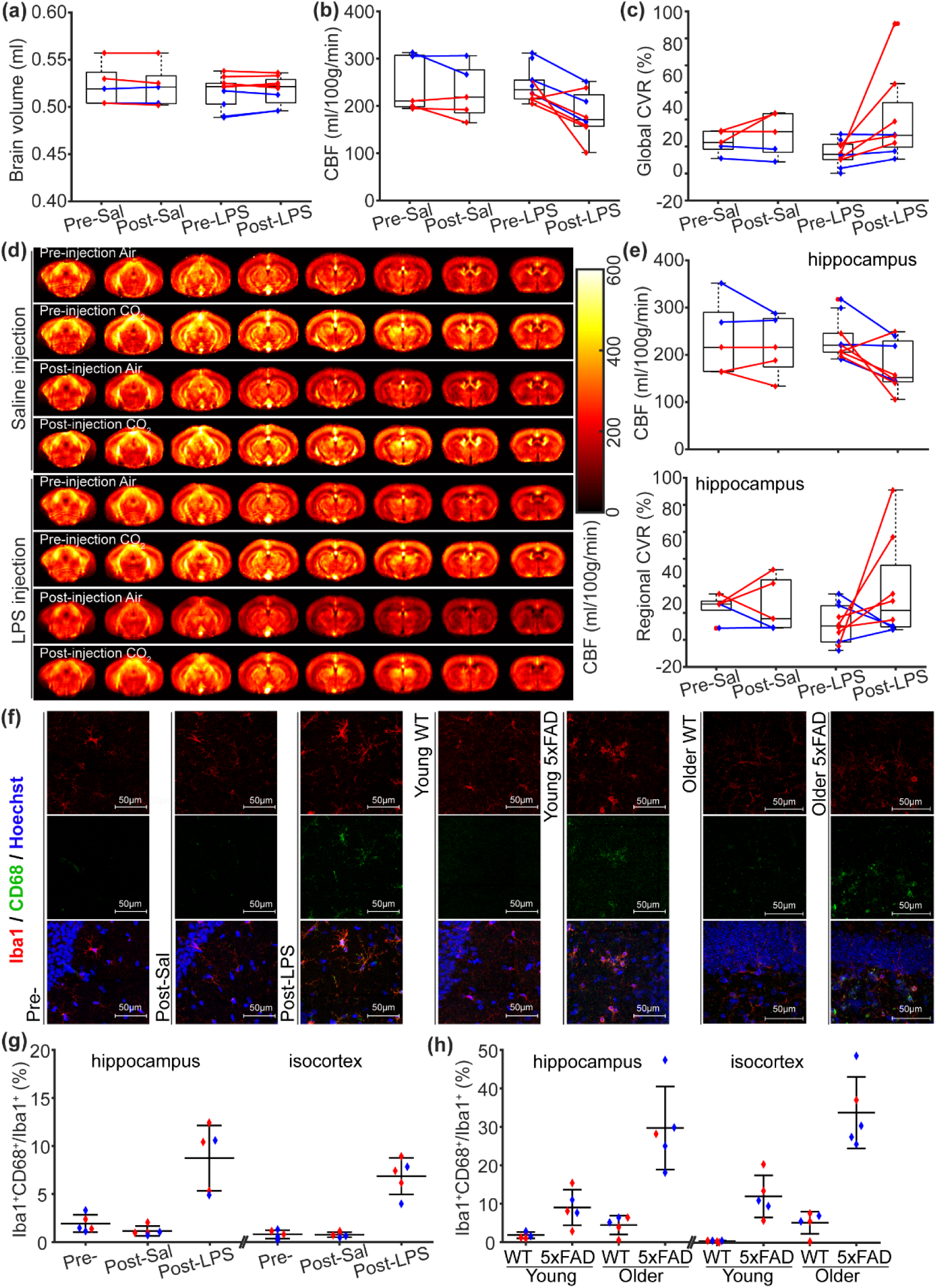
Characterization of the lipopolysaccharide (LPS)-induced neuroinflammation model. Comparisons of MRI measurements across pre-saline (Pre-Sal), post-saline (Post-Sal), pre-LPS, and post-LPS groups, including brain volume (a), cerebral blood flow (CBF; b), global cerebrovascular reactivity (CVR; c), regional mean CBF maps (d), and hippocampal CBF and CVR (e). Red and blue dots denote female and male mice, respectively. Dots connected by a line correspond to the same mouse. (f) Representative immunofluorescent images showing ionized calcium-binding adaptor molecule 1 (Iba1, red) and CD68 (green) staining in the hippocampus. Scale bars: 50 µm. Both the LPS-induced neuroinflammation and 5xFAD models are shown for comparison. (g) Comparisons of the CD68 coverage index across pre- (Pre-Sal and Pre-LPS; *n* = 5, 2F/3M), post-Sal (*n* = 5, 3F/2M), and post-LPS (*n* = 5, 3F/2M) groups in the hippocampus and isocortex. Error bars indicate standard deviation. The CD68 coverage index is defined as the ratio of the co-localized Iba1- and CD68-positive area (Iba1^+^CD68^+^) to the total Iba1-positive area (Iba1^+^). (h) Comparisons of the CD68 coverage index across young WT (*n* = 5, 3F/2M), young 5xFAD (*n* = 5, 3F/2M), older WT (*n* = 5, 3F/2M), and older 5xFAD (*n* = 5, 1F/4M) mice.

Immunofluorescent staining for Iba1 and CD68 was performed to assess the neuroinflammatory response. Representative high-magnification images demonstrated pronounced microglial activation after LPS injection, as well as in 5xFAD mice, within the hippocampus (Fig. 5f) and isocortex (Fig. S6). The CD68 coverage index, defined as the ratio of co-localized Iba1-positive and CD68-positive area (Iba1^+^CD68^+^) to the total Iba1-positive area (Iba1^+^), was quantified as a marker of inflammatory severity. Both the isocortex (*p* < 0.001; Fig. 5g) and hippocampus (*p* < 0.001; Fig. 5g) exhibited markedly elevated neuroinflammation after LPS injection. Co-staining for α-SMA and Collagen IV in the neuroinflammation model revealed no significant change in α-SMA coverage within either the hippocampus or isocortex after LPS treatment (Fig. S7). In 5xFAD mice, we repeated the co-staining of Iba1 and CD68 and found significant genotype × age interaction effects for the CD68 coverage index in both the hippocampus (*p* = 0.004; Fig. 5h) and isocortex (*p* = 0.002; Fig. 5h), confirming increased neuroinflammation.

Based on these results, the LPS injection induced robust microglial activation and reductions in baseline CBF within deep-brain regions, while CVR and α-SMA coverage remained preserved. These findings indicate that neuroinflammation primarily alters baseline CBF without impairing CVR.

## 4. DISCUSSION

In this study, we systematically characterized CVR in mouse models representing amyloidosis (5xFAD), VSMC loss (CADASIL), and neuroinflammation using non-contrast MRI techniques, along with corresponding histopathological assessments of α-SMA coverage, amyloid burden, and microglial activation. These models were chosen to disentangle the relative contributions of amyloid deposition, vascular structural integrity, and inflammation to cerebrovascular dysfunction—three major pathological components implicated in AD and related small-vessel disorders. We found that: (a) Aβ deposition occurred earlier in the pathological cascade than VSMC loss and CVR impairment; CVR impairment emerged only in the presence of VSMC loss, suggesting an association between VSMC degeneration and impaired vascular reactivity; (b) developmental VSMC deficiency in CADASIL mice led to marked CVR impairment, providing mechanistic evidence that VSMC loss is sufficient to impair CVR; and (c) neuroinflammation reduced baseline CBF without affecting CVR or α-SMA integrity, indicating a dissociation between inflammatory processes and CVR impairment. Collectively, these findings establish that CVR impairment is not a generic manifestation of neurodegeneration or inflammation, but rather a consequence of vascular contractile dysfunction arising from VSMC loss. In other words, VSMC loss—but not neuroinflammation—drives CVR impairment in AD.

The association between VSMC loss and CVR impairment reflects a disruption of vascular contractile function that compromises the vascular responses of small arteries and arterioles to metabolic or gas challenges. VSMCs provide the primary contractile force regulating arterial tone,^42^ and their loss would limit active vasoconstriction or vasodilation, thus flattening the cerebrovascular response to stimuli. This mechanism is supported by the CADASIL model, where developmental VSMC deficiency leads to CVR impairment despite preserved parenchymal integrity, confirming that vascular contractile dysfunction alone is sufficient to impair hemodynamic reactivity. In the pathological cascade of 5xFAD model, VSMC degeneration occurred secondarily to amyloid accumulation, indicating that amyloidosis may indirectly impair vascular responsiveness by damaging mural cells rather than by occluding or stiffening the lumen directly. Such VSMC loss could affect vascular compliance and impair signal propagation along the vascular tree,^43^ further restraining coordinated vascular dilation during hypercapnic or metabolic stimuli.

Our current preclinical findings parallel observations from human studies. In AD patients, impaired CO2-evoked or task-evoked vascular responses have been consistently reported using blood-oxygenation-level-dependent or arterial spin labeling MRI.^13,44-46^ Similarly, CADASIL patients exhibit blunted vasodilatory responses and delayed hemodynamic recovery,^47,48^ consistent with VSMC degeneration. The convergence between mouse and human data underscores that VSMC integrity is an important determinant of vascular reactivity across species and pathologies. Importantly, the preservation of CVR under inflammatory conditions in the LPS model suggests that neuroinflammation alone may not directly impair vasoreactivity, although it can modulate baseline perfusion and perivascular signaling^49^. These cross-model and cross-species consistencies reinforce the concept that cerebrovascular contractile dysfunction, rather than amyloid deposition or inflammation, represents an important driver of CVR impairment in AD.

Previous studies have primarily attributed vascular dysfunctions in AD to CAA, where vascular Aβ deposition stiffens vessel walls, disrupts endothelial function, and constrains vasodilation.^21,50,51^ CAA severity correlates with reduced vascular responsiveness, suggesting a potential contributing role in hemodynamic dysregulation. In these investigations, transgenic mouse models such as APP23, characterized by substantial CAA, were employed.^20^ The present study focuses on the 5xFAD mouse model that develops extensive parenchymal but only marginal vascular Aβ deposition. Despite minimal CAA involvement, 5xFAD mice exhibited CVR impairment coinciding with VSMC loss, suggesting that mural cell degeneration, rather than amyloid deposition within vessel walls, underlies the observed functional deficit. In other words, vascular contractile dysfunction can arise through VSMC loss even in the absence of substantial vascular amyloid accumulation, representing an alternative pathway to vascular dysregulation in AD. Platelet-derived growth factor (PDGF)-BB has been reported to mediate arterial stiffness^52^ and exhibits several-fold increases in both patients with mild cognitive impairment and 5xFAD mice.^53^ These findings suggest that PDGF-BB alteration may represent an upstream event in the CAA-independent pathway. Moreover, the dissociation from CAA-centered mechanisms underscores the importance of maintaining vascular smooth muscle cell (VSMC) integrity as an independent determinant of cerebrovascular health in the context of amyloid pathology.

The identification of VSMC loss as a critical contributor to CVR impairment has important translational implications. Since VSMCs are the principal mediators of vascular contractility,^42^ preserving their structural and functional integrity may represent a therapeutic avenue to maintain cerebrovascular responsiveness in AD. Agents targeting mural cell survival,^54^ smooth muscle contractility,^55^ or vascular remodeling^56^ could therefore mitigate hemodynamic dysfunction even in the absence of direct amyloid clearance. Moreover, MRI-based CVR mapping provides a noninvasive, non-contrast functional biomarker for detecting early vascular contractile deficits before overt neurodegeneration,^5^ and has been broadly used in clinical studies of AD, small-vessel disease, or aging.^12,13,16^ Integrating CVR assessment with other vascular imaging markers, such as blood-brain barrier permeability,^57^ may enable comprehensive evaluation of microvascular health in clinical settings. Current findings suggest that restoring vascular contractile competence may complement amyloid-targeted therapy^18^ in improving cerebrovascular resilience in AD.

Several limitations of this study should be acknowledged. First, although the current work establishes a strong temporal and mechanistic association between VSMC loss and CVR impairment, causality was inferred indirectly through comparative modeling rather than by selective genetic or pharmacological manipulation on VSMC. Future studies using inducible VSMC depletion or rescue models will be required to confirm the reversibility of this relationship. Second, the MRI resolution, while optimized for regional assessment, may have limited sensitivity to microregional heterogeneity in perfusion or reactivity within small vessels. Finally, the preclinical models examined here represent discrete pathological components—amyloidosis, VSMC loss, and inflammation—rather than the full spectrum of mixed pathologies^58,59^ in human AD. Despite these limitations, the comparative approach adopted here helps disentangle potential vascular mechanisms underlying CVR impairment.

In summary, this study delineates a mechanistic hierarchy of vascular dysfunction in AD, demonstrating that VSMC loss rather than amyloid deposition or neuroinflammation drives CVR impairment. Our results underscore that vascular contractile failure, rather than vascular amyloidosis or inflammatory tone, represents a critical determinant of hemodynamic dysfunction in AD. This work establishes a promising conceptual framework linking mural cell degeneration to cerebrovascular dysregulation and highlights the importance of preserving vascular integrity in neurodegenerative disease.

## ACKNOWLEDGMENTS

This work was supported by the Grant Sponsors: National Institutes of Health (NIH) R01 AG081932 and NIH P41 EB031771. The authors thank Dr. Angeliki Louvi (Yale University) for assistance in breeding the CADASIL mouse model and thank Dr. LaToya Ann Roker and Dr. Steven Wilbert (The Johns Hopkins University School of Medicine Microscope Facility) for assistance in microscope scanning.

## SUPPORTING INFORMATION

Additional supporting information can be found online in the Supporting Information section at the end of this article.

