## Supplemental Figures for "Vascular smooth muscle cell loss, but not neuroinflammation, drives cerebrovascular reactivity impairment in Alzheimer’s disease"

### Supporting Information

### 1. Cerebral blood flow (CBF) and cerebrovascular reactivity (CVR) in 5xFAD mice

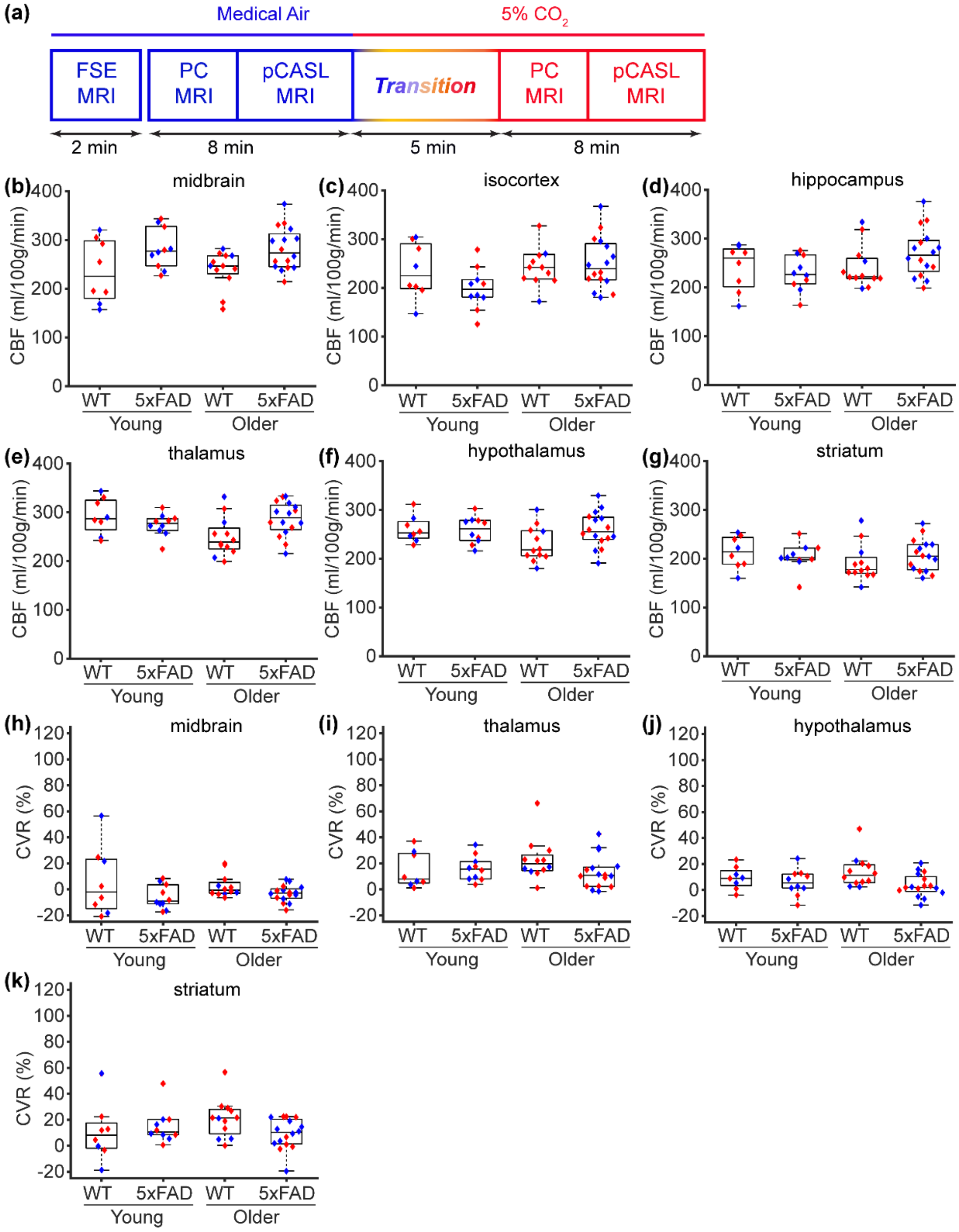

**Figure S1** Regional cerebral blood flow (CBF) and cerebrovascular reactivity (CVR) values in young and older 5xFAD mice. (a) Schematic diagram of the experimental design. Under medical air, a fast spin-echo (FSE) MRI sequence was used to estimate brain volume, and both phase-contrast (PC) and pseudo-continuous arterial spin labeling (pCASL) MRI were performed to measure baseline CBF. After switching to 5% CO2 gas, a 5-min transition period was used, after which PC and pCASL scans were repeated to assess regional CVR. (b–g) Regional CBF values in the midbrain, isocortex, hippocampus, thalamus, hypothalamus, and striatum, respectively. (h–k) Regional CVR values in the midbrain, thalamus, hypothalamus, and striatum, respectively.

### 2. Immunofluorescent images of α-smooth muscle actin (α-SMA) and Collagen IV staining in 5xFAD mice

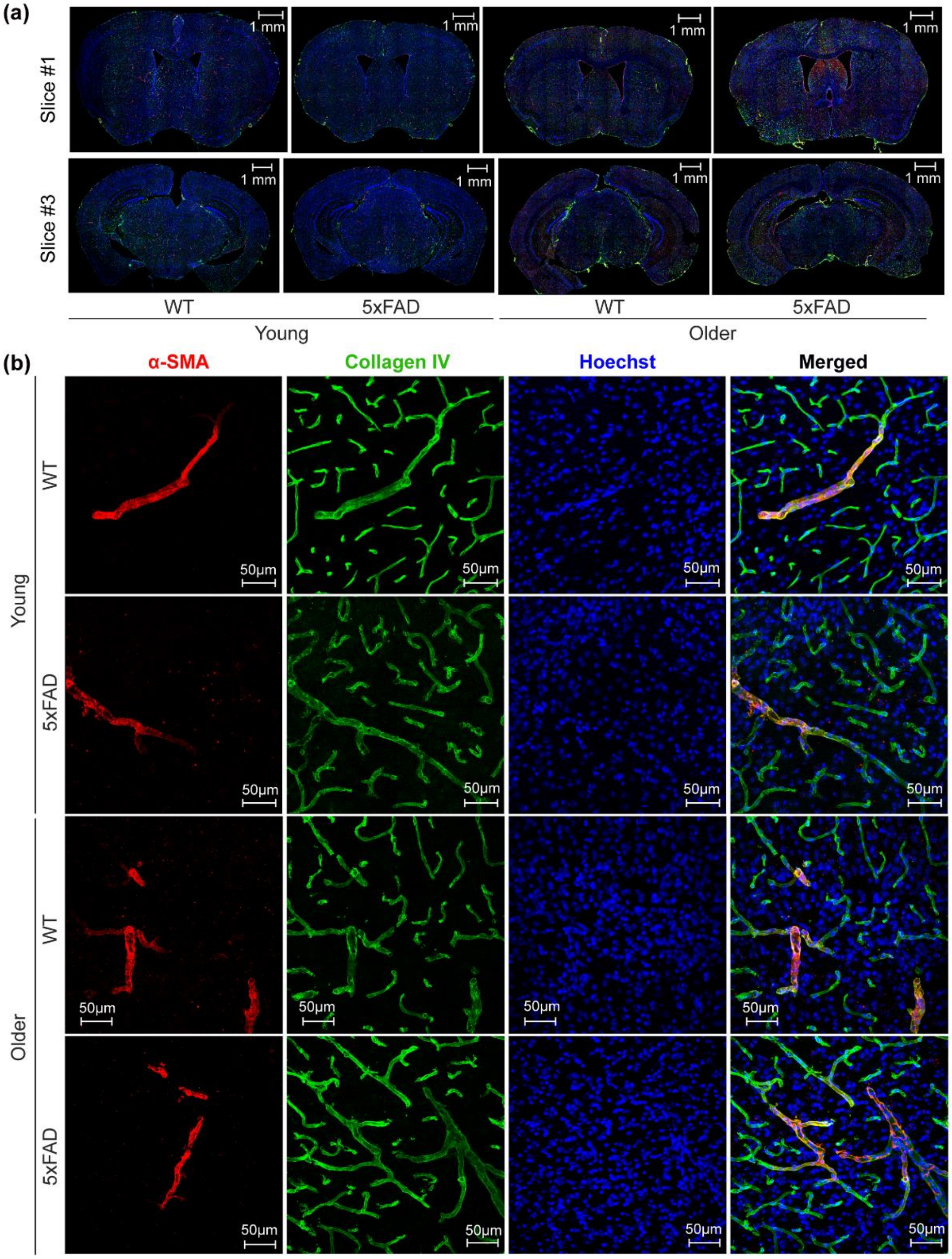


**Figure S2** (a) Whole-slice and (b) isocortical immunofluorescent images showing α-SMA (red) and Collagen IV (green) staining. Hoechst staining (blue) was used to visualize nuclei as a reference. Slice #1 to #3 indicate the rostral-to-caudal direction. Scale bars: 1 mm (whole slice) and 50 μm (isocortex).

**3. Regional CBF and CVR in cerebral autosomal dominant arteriopathy with subcortical infarcts and leukoencephalopathy (CADASIL) mice**

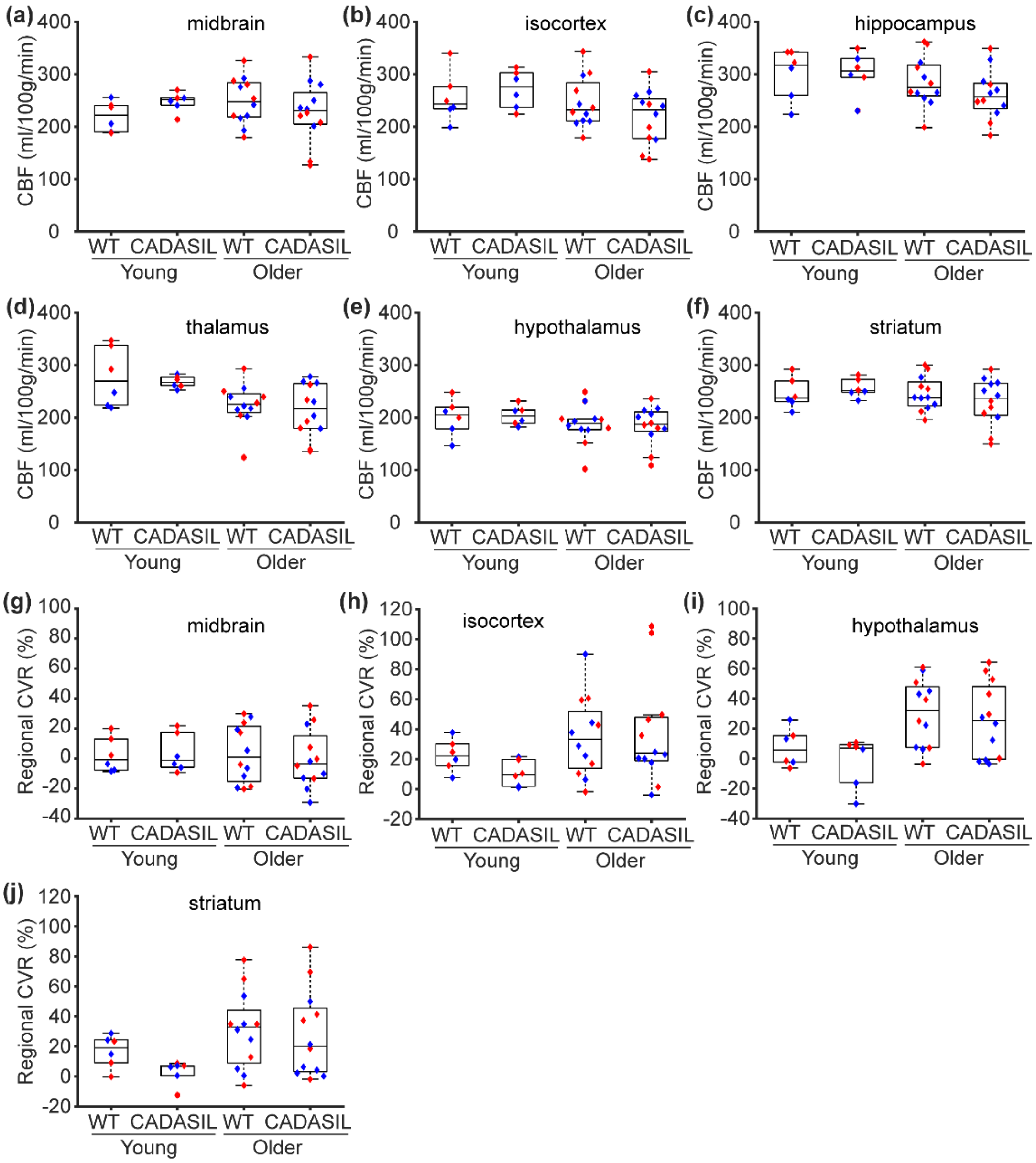


**Figure S3** Regional cerebral blood flow (CBF) and cerebrovascular reactivity (CVR) values in young and older cerebral autosomal dominant arteriopathy with subcortical infarcts and leukoencephalopathy (CADASIL) mice. (a–f) Regional CBF values in the midbrain, isocortex, hippocampus, thalamus, hypothalamus, and striatum, respectively. (g–j) Regional CVR values in the midbrain, isocortex, hypothalamus, and striatum, respectively.

**4. Immunofluorescent images of α-SMA and Collagen IV staining in the thalamus region of CADASIL mice**

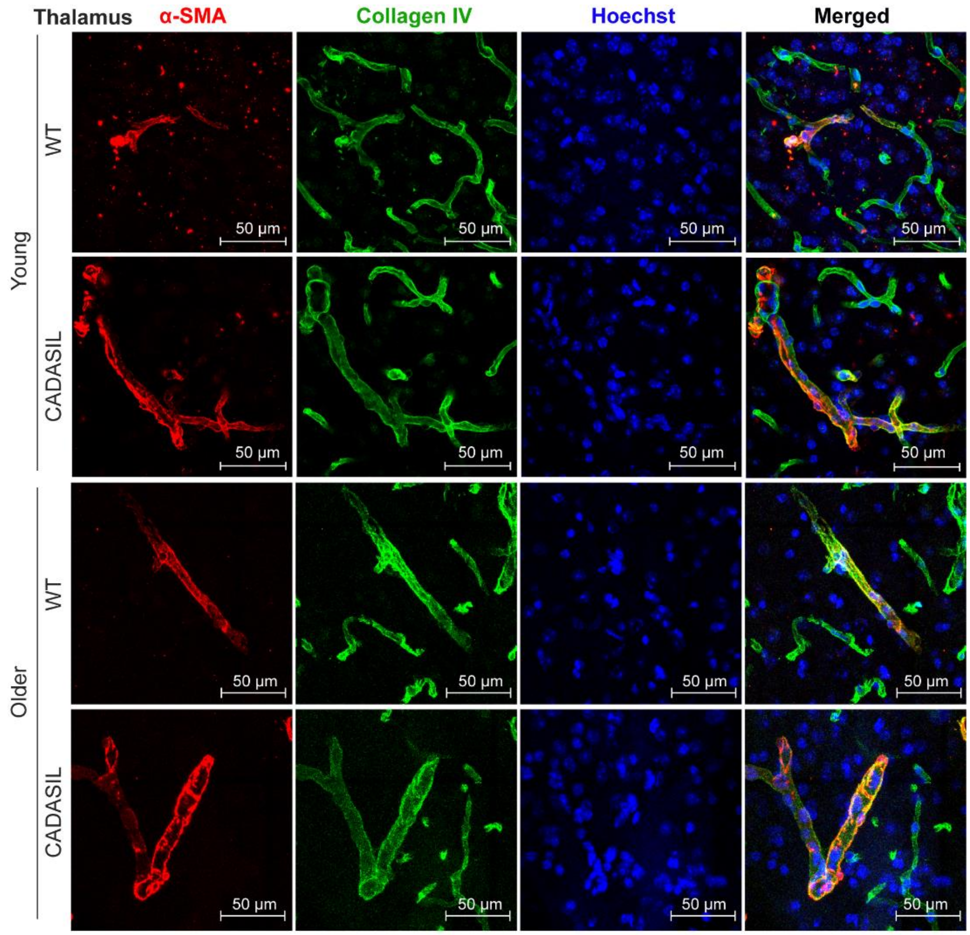


**Figure S4** Immunofluorescent images of α-smooth muscle actin (α-SMA; red) and Collagen IV (green) staining in the thalamus of cerebral autosomal dominant arteriopathy with subcortical infarcts and leukoencephalopathy (CADASIL) mice. Hoechst staining (blue) was used to visualize nuclei. Scale bar: 50 μm.

**5. Regional CBF and CVR in lipopolysaccharide (LPS)-induced neuroinflammation model**

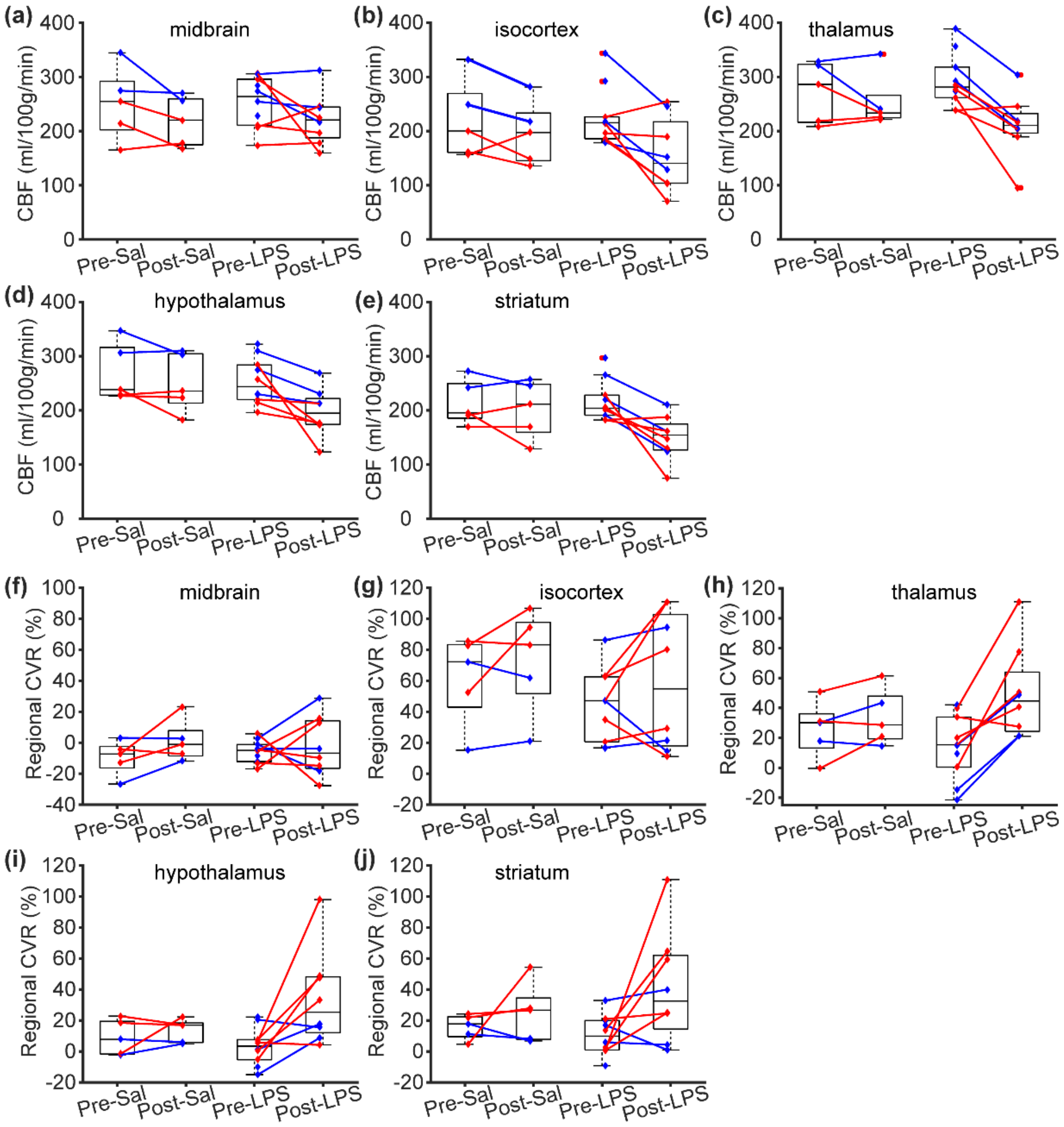


**Figure S5** Regional cerebral blood flow (CBF) and cerebrovascular reactivity (CVR) values before and after saline or lipopolysaccharide (LPS) injection. (a–e) Regional CBF values in the midbrain, isocortex, thalamus, hypothalamus, and striatum at pre-saline (Pre-Sal), post-saline (Post-Sal), pre-LPS, and post-LPS conditions. (f–j) Corresponding regional CVR values in the midbrain, isocortex, thalamus, hypothalamus, and striatum.

**6. Immunofluorescent images of ionized calcium-binding adaptor molecule 1 (Iba1) and CD68 staining in the neuroinflammation and 5xFAD models**

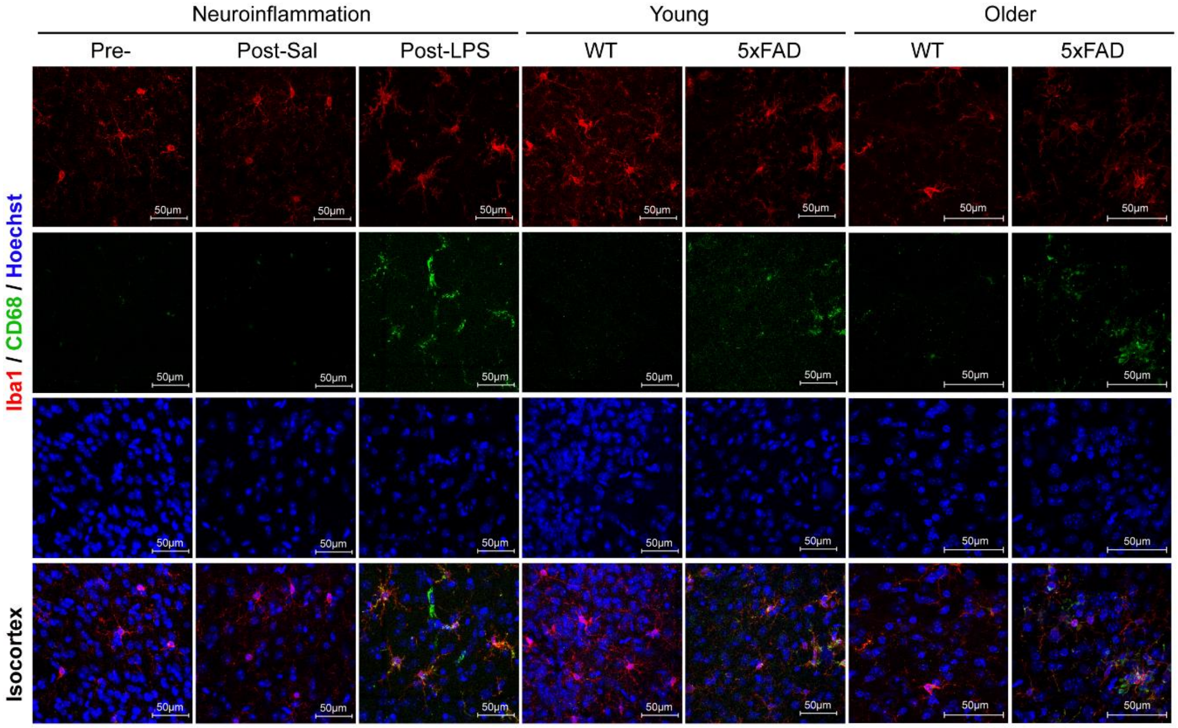


**Figure S6** Representative immunofluorescent images showing ionized calcium-binding adaptor molecule 1 (Iba1; red) and CD68 (green) staining in the isocortex of lipopolysaccharide (LPS)-induced neuroinflammation and 5xFAD mice. Hoechst staining (blue) was used to visualize nuclei. Scale bar: 50 μm.

### 7. Immunofluorescent images of α-SMA and Collagen IV staining in the neuroinflammation model

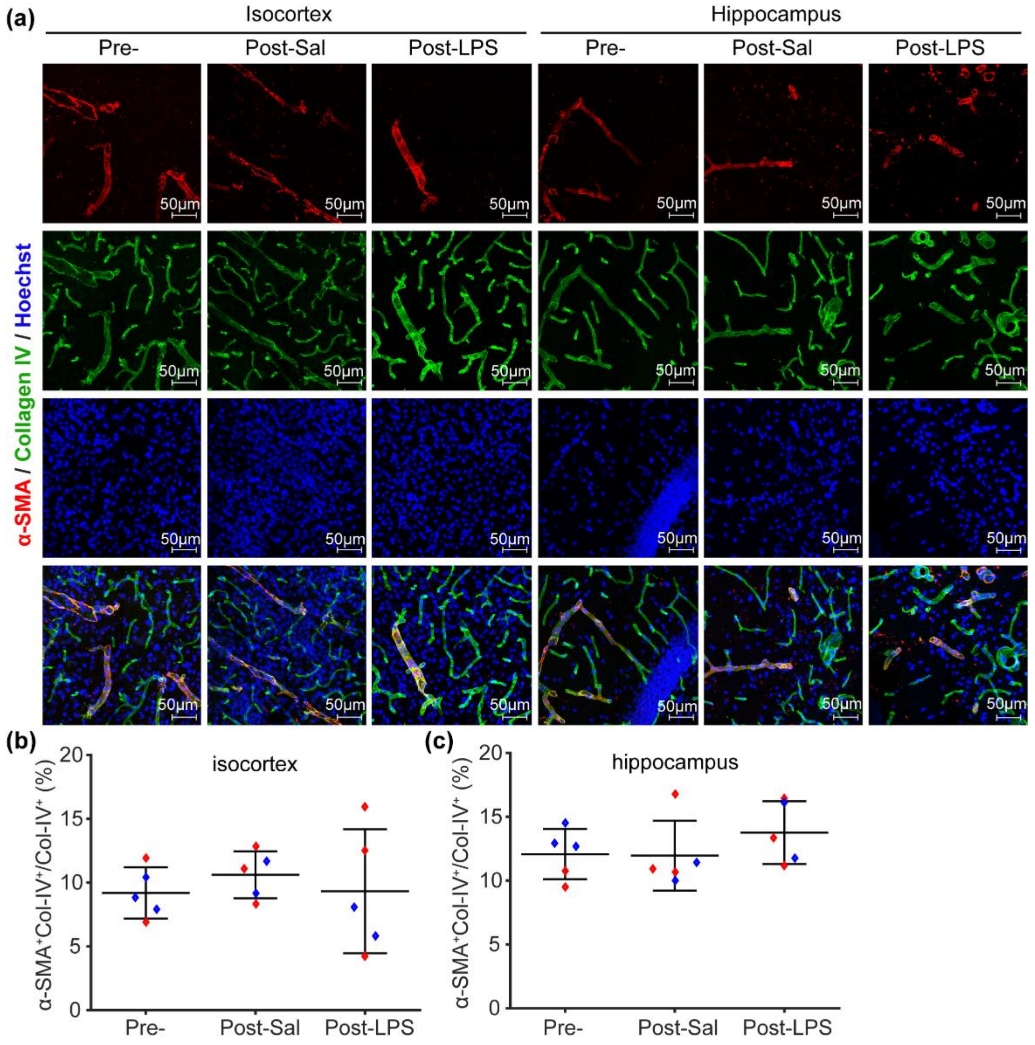


**Figure S7** α-smooth muscle actin (α-SMA) coverage in the neuroinflammation model. (a) Representative immunofluorescent images showing α-smooth muscle actin (α-SMA; red) and Collagen IV (Col-IV; green) staining under pre-injection (Pre-), post-saline (Post-Sal), and post-lipopolysaccharide (Post-LPS) conditions in the isocortex and hippocampus. Hoechst staining (blue) was used to visualize nuclei. Scale bar: 50 μm. (b, c) α-SMA coverage index in the isocortex and hippocampus, respectively, across the examined conditions. The α-SMA coverage index is defined as the ratio of the co-localized α-SMA– and Collagen IV–positive area (α-$SMA^+$Col-$IV^+$) to the total Collagen IV–positive area (Col-$IV^+$). Red and blue dots represent female and male mice, respectively. Error bars indicate standard deviation.

### 8. Detailed information of antibodies used in this study (Table S1)

| # | Name | Vendor | Catalog # | Host | Feature | Conc. (µg/mL) | Pack volume(µL) | Optimal Dilution | Storage (˚C) |
|---|---|---|---|---|---|---|---|---|---|
| 1 | Collagen IV | Bio-Rad | 2150-1470 | Rabbit | Poly | - | 100 | 1/500 | -20 |
| 2 | α-SMA | Sigma-Aldrich | C6198-.2ML | Mouse | Mono | - | 200 | 1/250 | 4 |
| 3 | Iba1 | FujiFilm | 019-19741 | Rabbit | - | 500-700 | 50 | 1/500 | -20 |
| 4 | CD68 | Bio-Rad | MCA1957 | Rat | Mono | 1000 | 250 | 1/250 | -20 |
| 5 | 6E10 | Biolegend | 803001 | Mouse | Mono | 1000 | 200 | 1/500 | 4 |

### 9. Details of statistical comparisons in this study (Table S2)

| Comparison | Statistical information | *p*-value | Statistical information | *p*-value | Statistical information | *p*-value |
|---|---|---|---|---|---|---|
| ***Session 1. Vascular and metabolic responses to hypercapnia*** | | | | | | |
| Student's *t*-test: $CBF_{air}$ vs. $CBF_{CO2}$ | *t*[11] = -4.547 | 8.346E-04 | | | | |
| *t*-test: $OEF_{air}$ vs. $OEF_{CO2}$ | *t*[11] = 2.665 | 0.022 | | | | |
| *t*-test: $CMRO2_{air}$ vs. $CMRO2_{CO2}$ | *t*[11] = -0.176 | 0.864 | | | | |
| Pearson correlation: CBF vs. CVR | $R^2$ = 0.740 | 2.128E-17 | | | | |
| ***Session 2. Characterization of 5xFAD model***[a] | | | | | | |
| Brain volume ~ genotype + age + sex | $\beta_{genotype}$=-1.76E-03 | 0.699 | $\beta_{age}$=5.65E-03 | 1.734E-08 | $\beta_{sex}$=-4.82E-06 | 0.999 |
| Global CBF ~ genotype + age + sex + rr | $\beta_{genotype}$=-3.43 | 0.745 | $\beta_{age}$=-1.03 | 0.579 | $\beta_{sex}$=-11.40 | 0.247 |
| CVR ~ genotype * age + sex + CBF | $\beta_{genotype*age}$=-2.68 | 0.036 | - | - | $\beta_{sex}$=0.67 | 0.841 |
| $CBF_{MB}$ ~ genotype + age + sex + rr | $\beta_{genotype}$=45.56 | 0.005 | $\beta_{age}$=-0.02 | 0.994 | $\beta_{sex}$=-4.50 | 0.756 |
| $CBF_{ISO}$ ~ genotype + age + sex + rr | $\beta_{genotype}$=-31.33 | 0.045 | $\beta_{age}$=2.30 | 0.392 | $\beta_{sex}$=-11.39 | 0.420 |
| $CBF_{HPC}$ ~ genotype + age + sex + rr | $\beta_{genotype}$=2.96 | 0.848 | $\beta_{age}$=1.40 | 0.607 | $\beta_{sex}$=-13.77 | 0.338 |
| $CBF_{TH}$ ~ genotype * age + sex + rr | $\beta_{genotype*age}$=8.83 | 0.022 | - | - | $\beta_{sex}$=-9.01 | 0.375 |
| $CBF_{HY}$ ~ genotype + age + sex + rr | $\beta_{genotype}$=2.90 | 0.781 | $\beta_{age}$=-4.51 | 0.018 | $\beta_{sex}$=-7.77 | 0.423 |
| $CBF_{STR}$ ~ genotype + age + sex + rr | $\beta_{genotype}$=-7.18 | 0.500 | $\beta_{age}$=-3.47 | 0.069 | $\beta_{sex}$=-9.07 | 0.358 |
| $CVR_{MB}$ ~ genotype + age + sex + $CBF_{MB}$ | $\beta_{genotype}$=-8.21 | 0.064 | $\beta_{age}$=0.17 | 0.828 | $\beta_{sex}$=-3.64 | 0.371 |
| $CVR_{ISO}$ ~ genotype * age + sex + $CBF_{ISO}$ | $\beta_{genotype*age}$=-6.53 | 0.027 | - | - | $\beta_{sex}$=-2.20 | 0.775 |
| $CVR_{HPC}$ ~ genotype * age + sex + $CBF_{HPC}$ | $\beta_{genotype*age}$=-3.29 | 0.033 | - | - | $\beta_{sex}$=-0.48 | 0.905 |
| $CVR_{TH}$ ~ genotype + age + sex + $CBF_{TH}$ | $\beta_{genotype}$=-6.80 | 0.134 | $\beta_{age}$=0.23 | 0.771 | $\beta_{sex}$=-1.45 | 0.729 |
| $CVR_{HY}$ ~ genotype + age + sex + $CBF_{HY}$ | $\beta_{genotype}$=-6.70 | 0.054 | $\beta_{age}$=0.45 | 0.460 | $\beta_{sex}$=1.27 | 0.691 |
| $CVR_{STR}$ ~ genotype + age + sex + $CBF_{STR}$ | $\beta_{genotype}$=-4.07 | 0.430 | $\beta_{age}$=0.36 | 0.697 | $\beta_{sex}$=5.20 | 0.282 |
| Young $CVR_{HPC}$ ~ genotype + sex + $CBF_{HPC}$ | $\beta_{genotype}$=4.64 | 0.598 | - | - | $\beta_{sex}$=-2.68 | 0.740 |
| Older $CVR_{HPC}$ ~ genotype + sex + $CBF_{HPC}$ | $\beta_{genotype}$=-17.05 | 3.064E-04 | - | - | $\beta_{sex}$= 2.04 | 0.607 |
| $CI(\alpha\text{-SMA/Col-IV})_{ISO}$ ~ genotype * age + sex | $\beta_{genotype*age}$=-0.20 | 0.023 | - | - | $\beta_{sex}$=0.20 | 0.843 |
| $CI(\alpha\text{-SMA/Col-IV})_{HPC}$ ~ genotype * age + sex | $\beta_{genotype*age}$=-0.23 | 0.024 | - | - | $\beta_{sex}$=-0.88 | 0.440 |
| *t*-test: Young WT vs. 5xFAD CI(α-SMA/Col- | *t*[8] = 1.833 | 0.104 | | | | |

| | | | | | | |
|---|---|---|---|---|---|---|
| IV)ISO | | | | | | |
| *t*-test: Older WT vs. 5xFAD CI(α-SMA/Col-IV)ISO | *t*[8] = 4.263 | 0.003 | | | | |
| *t*-test: Young WT vs. 5xFAD CI(α-SMA/Col-IV)HPC | *t*[8] = 0.731 | 0.485 | | | | |
| *t*-test: Older WT vs. 5xFAD CI(α-SMA/Col-IV)HPC | *t*[8] = 3.213 | 0.012 | | | | |
| CI(6E10/Region)ISO ~ genotype * age + sex | $\beta_{genotype*age}$=0.07 | 0.001 | - | - | $\beta_{sex}$=-0.19 | 0.415 |
| CI(6E10/Region)HPC ~ genotype * age + sex | $\beta_{genotype*age}$=0.10 | 6.710E-04 | - | - | $\beta_{sex}$=-0.36 | 0.234 |
| *t*-test: Young WT vs. 5xFAD CI(6E10/Region)ISO | *t*[8] = -9.16 | 1.625E-05 | | | | |
| *t*-test: Older WT vs. 5xFAD CI(6E10/Region)ISO | *t*[8] = -5.75 | 4.274E-04 | | | | |
| *t*-test: Young WT vs. 5xFAD CI(6E10/Region)HPC | *t*[8] = -7.41 | 7.516E-05 | | | | |
| *t*-test: Older WT vs. 5xFAD CI(6E10/Region)HPC | *t*[8] = -5.99 | 3.281E-04 | | | | |
| *t*-test: young vs. older CI(Col-IV/6E10)ISO | *t*[8] = -2.677 | 0.028 | | | | |
| *t*-test: young vs. older CI(Col-IV/6E10)HPC | *t*[8] = -5.179 | 8.435E-04 | | | | |
| CI(CD68/Iba1)HPC ~ genotype * age + sex | $\beta_{genotype*age}$= 0.86 | 0.004 | - | - | $\beta_{sex}$=-2.92 | 0.351 |
| CI(CD68/Iba1)ISO ~ genotype * age + sex | $\beta_{genotype*age}$= 0.90 | 0.002 | - | - | $\beta_{sex}$=-0.74 | 0.794 |
| ***Session 3. Characterization of CADASIL model*** | | | | | | |
| Brain volume ~ genotype + age + sex | $\beta_{genotype}$=-3.82E-03 | 0.498 | $\beta_{age}$=-1.40E-03 | 0.255 | $\beta_{sex}$=0.02 | 0.010 |
| Global CBF ~ genotype + age + sex + rr | $\beta_{genotype}$=-4.03 | 0.744 | $\beta_{age}$=-5.94 | 0.169 | $\beta_{sex}$=0.23 | 0.985 |
| CVR ~ genotype * age + sex + CBF | $\beta_{genotype}$=-6.49 | 0.024 | $\beta_{age}$=1.88 | 0.004 | $\beta_{sex}$=-7.87 | 0.007 |
| $CBF_{MB}$ ~ genotype + age + sex + rr | $\beta_{genotype}$=-4.43 | 0.787 | $\beta_{age}$=0.73 | 0.897 | $\beta_{sex}$=6.94 | 0.668 |
| $CBF_{ISO}$ ~ genotype + age + sex + rr | $\beta_{genotype}$=-10.18 | 0.553 | $\beta_{age}$=-8.89 | 0.139 | $\beta_{sex}$=-8.61 | 0.610 |
| $CBF_{HPC}$ ~ genotype + age + sex + rr | $\beta_{genotype}$=-11.41 | 0.476 | $\beta_{age}$=-9.36 | 0.096 | $\beta_{sex}$=-15.62 | 0.324 |
| $CBF_{TH}$ ~ genotype * age + sex + rr | $\beta_{genotype}$=-5.08 | 0.741 | $\beta_{age}$=-15.22 | 0.007 | $\beta_{sex}$=4.60 | 0.761 |
| $CBF_{HY}$ ~ genotype + age + sex + rr | $\beta_{genotype}$=1.96 | 0.870 | $\beta_{age}$=-5.72 | 0.174 | $\beta_{sex}$=5.97 | 0.615 |
| $CBF_{STR}$ ~ genotype + age + sex + rr | $\beta_{genotype}$=-4.03 | 0.744 | $\beta_{age}$=-5.94 | 0.169 | $\beta_{sex}$=0.23 | 0.985 |
| $CVR_{MB}$ ~ genotype + age + sex + $CBF_{MB}$ | $\beta_{genotype}$=-3.19 | 0.552 | $\beta_{age}$=-0.47 | 0.686 | $\beta_{sex}$=-11.77 | 0.033 |

| | | | | | | |
|---|---|---|---|---|---|---|
| $CVR_{ISO}$ ~ genotype + age + sex + $CBF_{ISO}$ | $\beta_{genotype}$=-6.85 | 0.191 | $\beta_{age}$=2.58 | 0.028 | $\beta_{sex}$=-14.30 | 0.009 |
| $CVR_{HPC}$ ~ genotype + age + sex + $CBF_{HPC}$ | $\beta_{genotype}$=-4.32 | 0.226 | $\beta_{age}$=1.85 | 0.022 | $\beta_{sex}$=-7.40 | 0.042 |
| $CVR_{TH}$ ~ genotype + age + sex + $CBF_{TH}$ | $\beta_{genotype}$=-9.75 | 0.026 | $\beta_{age}$=3.01 | 0.002 | $\beta_{sex}$=0.37 | 0.929 |
| $CVR_{HY}$ ~ genotype + age + sex + $CBF_{HY}$ | $\beta_{genotype}$=-9.93 | 0.028 | $\beta_{age}$=3.94 | 2.245E-04 | $\beta_{sex}$=-14.51 | 0.002 |
| $CVR_{STR}$ ~ genotype + age + sex + $CBF_{STR}$ | $\beta_{genotype}$=-10.49 | 0.030 | $\beta_{age}$=2.59 | 0.016 | $\beta_{sex}$=-12.29 | 0.012 |
| $CI(\alpha\text{-SMA/Col-IV})_{HPC}$ ~ genotype + age + sex | $\beta_{genotype}$=-1.71 | 0.113 | $\beta_{age}$=0.14 | 0.492 | $\beta_{sex}$=-1.66 | 0.124 |
| $CI(\alpha\text{-SMA/Col-IV})_{TH}$ ~ genotype + age + sex | $\beta_{genotype}$=-2.01 | 0.008 | $\beta_{age}$=-0.17 | 0.229 | $\beta_{sex}$=0.14 | 0.833 |
| ***Session 4. Characterization of neuroinflammation model***[b] | | | | | | |
| Brain volume ~ injection + sex + ε | $\beta_{saline}$=2.13E-03 | 0.775 | $\beta_{LPS}$=-2.47E-03 | 0.696 | $\beta_{sex}$=0.02 | 4.669E-04 |
| Global CBF ~injection + sex + rr + ε | $\beta_{saline}$=-13.53 | 0.439 | $\beta_{LPS}$=-39.46 | 0.039 | $\beta_{sex}$=-62.75 | 5.464E-05 |
| Global CVR ~ injection + sex + CBF + ε | $\beta_{saline}$=7.68 | 0.381 | $\beta_{LPS}$=6.88 | 0.459 | $\beta_{sex}$=1.73 | 0.848 |
| $CBF_{MB}$ ~ injection + sex + rr + ε | $\beta_{saline}$=-31.29 | 0.156 | $\beta_{LPS}$=-21.05 | 0.357 | $\beta_{sex}$=-56.84 | 0.001 |
| $CBF_{ISO}$ ~ injection + sex + rr + ε | $\beta_{saline}$=-25.79 | 0.350 | $\beta_{LPS}$=-45.16 | 0.126 | $\beta_{sex}$=-61.81 | 0.005 |
| $CBF_{HPC}$ ~ injection + sex + rr + ε | $\beta_{saline}$=-13.53 | 0.536 | $\beta_{LPS}$=-20.90 | 0.365 | $\beta_{sex}$=-62.34 | 6.895E-04 |
| $CBF_{TH}$ ~ injection + sex + rr + ε | $\beta_{saline}$=-31.24 | 0.152 | $\beta_{LPS}$=-56.66 | 0.018 | $\beta_{sex}$=-65.04 | 3.662E-04 |
| $CBF_{HY}$ ~ injection + sex + rr + ε | $\beta_{saline}$=-6.49 | 0.698 | $\beta_{LPS}$=-38.91 | 0.035 | $\beta_{sex}$=-63.12 | 3.024E-05 |
| $CBF_{STR}$ ~ injection + sex + rr + ε | $\beta_{saline}$=-13.15 | 0.457 | $\beta_{LPS}$=-46.79 | 0.017 | $\beta_{sex}$=-45.71 | 0.002 |
| $CVR_{MB}$ ~ injection + sex + $CBF_{MB}$ + ε | $\beta_{saline}$=8.34 | 0.257 | $\beta_{LPS}$=5.79 | 0.455 | $\beta_{sex}$=0.42 | 0.956 |
| $CVR_{ISO}$ ~ injection + sex + $CBF_{ISO}$ + ε | $\beta_{saline}$=17.09 | 0.252 | $\beta_{LPS}$=-18.30 | 0.250 | $\beta_{sex}$=-4.99 | 0.744 |
| $CVR_{HPC}$ ~ injection + sex + $CBF_{HPC}$ + ε | $\beta_{saline}$=5.09 | 0.637 | $\beta_{LPS}$=1.14 | 0.920 | $\beta_{sex}$=-3.47 | 0.757 |
| $CVR_{TH}$ ~ injection + sex + $CBF_{TH}$ + ε | $\beta_{saline}$=13.85 | 0.258 | $\beta_{LPS}$=25.97 | 0.053 | $\beta_{sex}$=12.17 | 0.337 |
| $CVR_{HY}$ ~ injection + sex + $CBF_{HY}$ + ε | $\beta_{saline}$=5.38 | 0.552 | $\beta_{LPS}$=17.81 | 0.074 | $\beta_{sex}$=1.97 | 0.833 |
| $CVR_{STR}$ ~ injection + sex + $CBF_{STR}$ + ε | $\beta_{saline}$=9.18 | 0.401 | $\beta_{LPS}$=15.97 | 0.176 | $\beta_{sex}$=2.26 | 0.841 |
| $CI(CD68/Iba1)_{HPC}$ ~ injection + sex | $\beta_{saline}$=-0.91 | 0.511 | $\beta_{LPS}$=6.67 | 4.437E-04 | $\beta_{sex}$=0.67 | 0.559 |
| $CI(CD68/Iba1)_{ISO}$ ~ injection + sex | $\beta_{saline}$=-0.19 | 0.794 | $\beta_{LPS}$=5.91 | 4.224E-06 | $\beta_{sex}$=0.76 | 0.215 |
| $CI(\alpha\text{-SMA/Col-IV})_{HPC}$ ~ injection + sex | $\beta_{saline}$=-0.02 | 0.992 | $\beta_{LPS}$=1.78 | 0.290 | $\beta_{sex}$=-0.47 | 0.726 |
| $CI(\alpha\text{-SMA/Col-IV})_{ISO}$ ~ injection + sex | $\beta_{saline}$=1.11 | 0.603 | $\beta_{LPS}$=-0.18 | 0.932 | $\beta_{sex}$=1.54 | 0.385 |

[a] rr: respiration rate; MB: midbrain; ISO: isocortex; HPC: hippocampus; TH: thalamus; HY: Hypothalamus; STR: striatum. $CI(\alpha\text{-SMA/Col-IV})_{ISO}$: Coverage index (CI) defined by the ratio between $\alpha$-SMA$^{+}$Col-IV$^{+}$ co-localization and Col-IV$^{+}$ for the ISO region. Similar definitions followed the identical principle.

[b] Linear mixed-effects (LME) models were used for this part when relevant because repeated measurements in same mice were involved. A random term ($\varepsilon$) was included for each mouse in statistical analyses.
